# Dissecting multi-gene virulence phenotypes with base editing requires consideration of target-independent mutagenesis

**DOI:** 10.64898/2026.02.11.705302

**Authors:** Joshua P. M. Newson, Sandra Gawlitt, Sneha Sundar, Roland R. Regoes, Chase L. Beisel, Wolf-Dietrich Hardt

## Abstract

Bacterial pathogenicity arises from complex genetic interactions that are difficult to characterise through single-gene deletions. CRISPR base editors can generate multiplexed gene knockouts, yet this technology remains unexplored for dissecting bacterial pathogenicity. Here, we developed a base-editing pipeline for multi-gene knockouts while revealing that target-independent editing can contribute to variability in clonal fitness. In the model pathogen *Salmonella* Typhimurium, we employed curable plasmids containing a cytidine deaminase base editor and a multi-spacer CRISPR array to introduce premature stop codons in up to nine genes encoding SPI-2 T3SS effector proteins. Target bases were efficiently edited, producing a multi-knockout strain that showed reduced virulence *in vivo* relative to single knockouts. However, whole-genome sequencing revealed off-target cytidine deaminase activity, which affected virulence *in vivo* in a clone-dependent manner. A statistical power analysis predicted how many edited mutants are needed to confidently measure fitness functions in the face of off-target mutations. Our work shows the potential and current limitations of multiplexed base editing in bacterial pathogens and highlights the need for properly addressing off-target mutations when deploying base editors to interrogate genotype-phenotype relationships.

## Introduction

Pathogenic bacteria rely on complex genetic networks that permit host colonisation and transmission in a manner that must be genetically co-ordinated, environmentally responsive, and evolutionarily adaptable (1, 2). For example, genetic networks underpin the translocation of bacterial effector proteins into host cells via the activity of type-three secretion systems (T3SS) (3, 4). Secretion systems are typically encoded in one locus, while effector genes are often dispersed over the entire genome, as has been reported for a range of clinically relevant species of *Salmonella* (5, 6), *Shigella* (7), *Yersinia* (8), as well as various pathotypes of *E.coli* (9). While natural selection favours robust networks that tolerate genetic plasticity and interdependency, scientific interrogation of single gene deletions is typically used to understand genetic-phenotypic relationships. This approach is confounded by the sheer number of bacterial effectors, which often act cooperatively or redundantly. Further, current tools for genetic engineering are low-throughput, laborious, and typically limited to deletion or disruption of one genetic region (10, 11). Future advances in understanding the complex genetic basis of host-pathogen interactions will require new technologies that might permit genetic engineering in a manner that is faster, scalable, and more efficient in time and resources.

A range of CRISPR-based technologies provide the means for efficiently disrupting multiple bacterial genes simultaneously (12, 13). A widely-used option is CRISPR interference (CRISPRi) which can silence gene expression by employing a catalytically dead Cas9 (dCas9) and a guide RNA that targets a complementary sequence flanked by a downstream protospacer adjacent motif (PAM) (14, 15). While this technology has been successful in silencing multiple genes in bacterial pathogens (16), there are various limitations including variation in degrees of gene silencing, and cytotoxicity due to the bad-seed effect arising from partial binding of the guide to promoters of essential genes (17, 18). Alternatively, catalytically active Cas9 combined with a guide RNA can be exploited to achieve permanent modification of a target DNA sequence (19, 20), though this approach is not reliable for multiplexed genetic engineering and has proven inefficient in a range of bacterial species (21, 22). CRISPR base editors represent a promising alternative, which more precisely catalyse the conversion of individual target bases without inducing double-stranded breaks in DNA (23, 24). Cytidine base editors comprise a cytidine deaminase domain fused to a dead or nicking Cas9, which catalyses targeted deamination of a cytosine within the editing window. Subsequent DNA replication or repair permanently converts the cytosine to a thymine, resulting in a point mutation (23). Exploiting this catalytic activity via rational guide design to target specific codons can be used to achieve gene disruption by introduction of a premature stop codon into a given DNA sequence (25, 26), or similarly by disruption of an existing start codon (27). Base editors of this class combined with single guide RNAs (sgRNA) have been used to disrupt single genes in a range of bacterial pathogens including *Klebsiella pneumoniae* (28)*, Shigella flexneri* (29), and *Staphylococcus aureus* (30). The use of multiple guide RNAs to achieve multiplexed editing has been useful in exploring bacterial metabolism (31) and natural product synthesis (32), but much work remains to establish the potential of this strategy for dissecting mechanisms of host-pathogen interactions.

Here, we applied cytidine deaminase base editors to simultaneously disrupt cohorts of *Salmonella* Typhimurium SPI-2 effectors, and assessed the virulence of these edited strains *in vivo.* We show that the combination of temperature-sensitive cytidine deaminase base editors with multiplexed CRISPR guide arrays can reproducibly achieve editing of three target genes simultaneously, and in a manner that is iterative to produce even more complex mutants. In mouse models of infection, a base-edited strain showed a strong virulence defect similar to a mutant strain constructed by traditional recombineering. However, we found that base editing reactions also produce a range of other mutations in bacterial chromosomes, which we attribute to cytidine deaminase off-target activity. By performing parallel editing reactions in isogenic strains bearing genomic tags compatible with RT-qPCR, we report a variable level of off-target editing that produces a range of virulence phenotypes *in vivo* that may confound accurate discovery of genetic interactions. To address this technical problem, we performed a statistical power analysis and determined the number of base-edited clones needed to accurately predict unknown phenotypes. This provides a robust approach to deal with off-target mutations with implications for the broader application of CRISPR array-guided base editors in the study of bacterial pathogenesis and other cellular processes.

## Results

### Simultaneous disruption of multiple genes in *Salmonella* using a cytidine deaminase base editor

The development of novel genetic engineering approaches that are programmable, multiplexed, and logistically simple should ideally be contrasted to existing technologies that have proven to be reliable and efficient, and for which model organisms are available to explore the application of these techniques. *Salmonella* Typhimurium (*S*.Tm) represents an ideal model species, given the ease of genetic manipulation and culture conditions, the availability of various *in vitro* and *in vivo* models of disease, and the wide range of literature that describes the function of virulence genes to the pathogenicity of this organism (5, 6, 33). Several reports describe how deletion of single effector genes can have seemingly little impact on virulence (**Fig. 1A**) (11, 34, 35), and the creation of more complicated bacterial mutants has been limited by existing genetic tools (10, 11). Thus, we chose *S.*Tm genes encoding bacterial effectors of the SPI-2 T3SS as targets to contrast existing methods with new technologies for genetic engineering.

**Figure 1.**
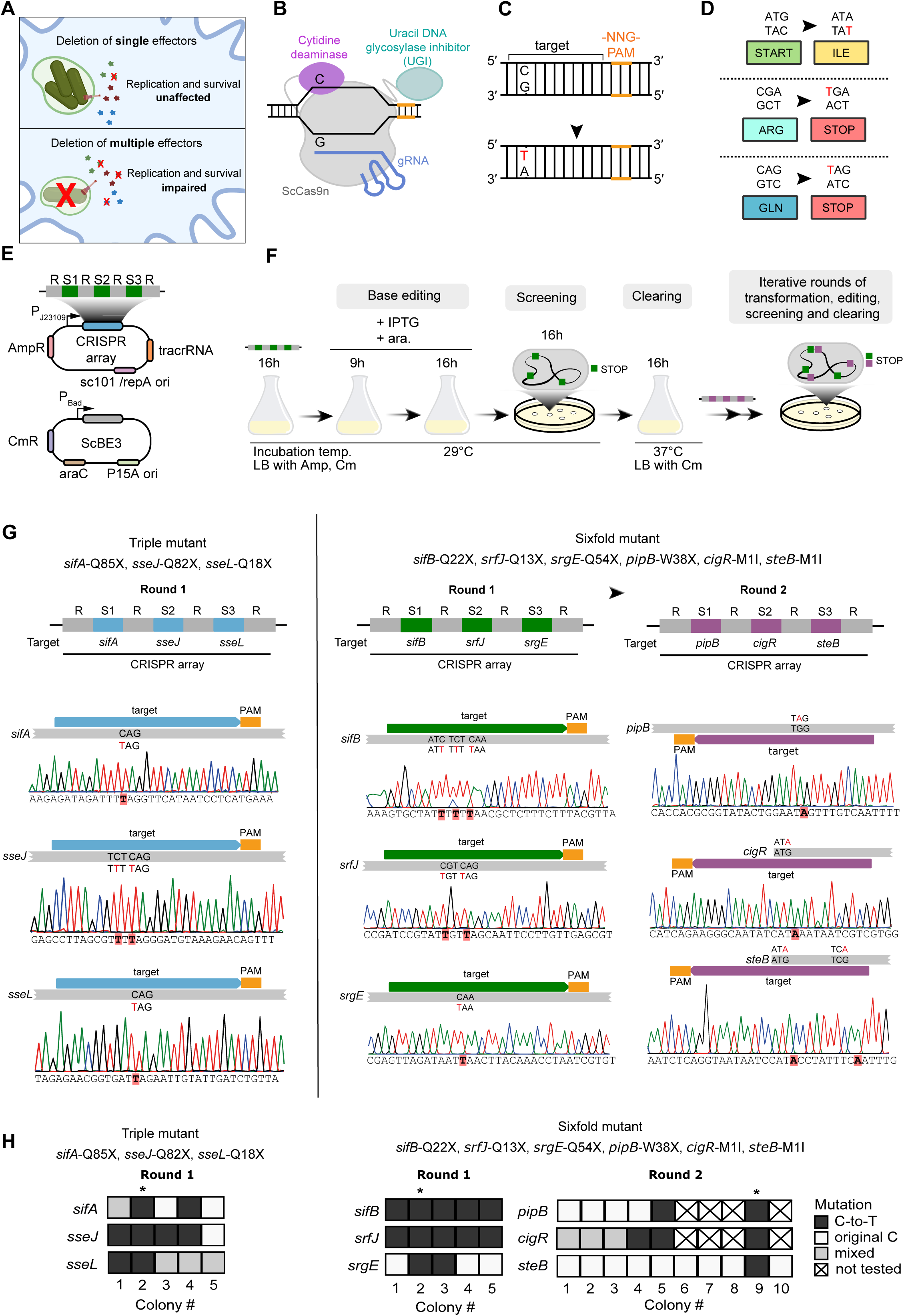
Multiplexed base-editing of target virulence genes in *Salmonella* Typhimurium using temperature-sensitive Cas9-CRISPR array plasmids. **A)** Cartoon depiction of the role of the SPI-2 type III secretion system (T3SS) in facilitating intracellular infection via translocation of bacterial effector proteins. Mutant strains deficient for single effector genes often phenocopy the wild-type parent strain, likely due to functional interdependency or redundancy between effectors. Deletion of multiple genes that are functionally related can result in strains that are attenuated for virulence. **B)** Scheme of the rAPO-ScCas9n-UGI (ScBE3) base editor, which was constructed by fusing ScCas9 D10A N-terminally to a rAPOBEC1 deaminase using a 16 amino-acid linker and fusing it C-terminally to a uracil DNA glycosylase inhibitor (UGI) using a 5 amino-acid linker. **C)** Scheme of a target locus with the target cytosine that is converted into a thymine through base editing. The protospacer adjacent motif (PAM) is depicted in orange, the mutated base in red. **D)** Nucleobase context of ScBE3 target-cytosines that lead to premature stop codons or disrupted start codons. Base-edited changes to target codon shown in red. **E)** Cartoon schematic of the two constructs used for multiplexed base editing of *Salmonella* effector genes. The temperature-sensitive CRISPR array plasmid (upper) encodes three constitutively expressed spacers targeting different genes and is maintained at temperatures below 30 °C. The base editor plasmid (lower) encodes ScBE3 under control of an inducible IPTG and L-arabinose hybrid promoter. **F)** Overview of the experimental procedure to generate a multi-effector knockout strain using sequential rounds of multiplexed base editing. *S.*Tm is first co-transformed with the base editor and a triple-spacer array plasmid (Fig. 1E). The transformants are then cultured for 16 hours in LB medium supplemented with antibiotics. The next day, the culture is diluted in LB medium supplemented with IPTG and L-arabinose to induce base editing and incubated for 9 hours. Another subculturing step for 16 hours increases editing efficiency. To maintain the temperature-sensitive array plasmid (AmpR), the culture is incubated at 29°C. After screening for successful editing, the confirmed clones are cultured in medium without ampicillin at 37°C to clear the cells from the CRISPR array plasmid. The procedure can be repeated with new CRISPR array plasmids until all gene targets have been disrupted by sequential rounds of base editing. **G)** Generation of two mutant strains of *S*.Tm, deficient for indicated effector genes via base editing of premature stop codons. A triple mutant (*sifA-*Q85X, *sseJ*-Q82X, *sseL*-Q18X) was generated in one round of editing (left, blue spacers), while a sixfold mutant (*sifB*-Q22X, *srfJ*-Q13X, *srgE*-Q54X, *pipB*-W38X, *cigR*-M1I, *steB*-M1I) was generated in two sequential rounds of editing (right, green and purple spacers). Sanger sequencing was used to confirm editing of target bases (red). **H)** Quantification of editing frequency based on Sanger sequencing of colony PCR amplicons. Successfully edited target bases shown in black, unedited target bases shown in white, ‘not tested’ indicates genes were not screened by colony PCR. Clones indicated with an asterisk (*) were selected for experimental use or further iterative editing, as required.

To enable rapid disruption of multiple genes, we employed a cytidine base editor comprising the ScCas9 nickase fused to APOBEC cytidine deaminase and a uracil DNA glycosylase inhibitor domain (termed ScBE3) (36). The base editor converts a cytosine within the editing window of the displaced strand into a uracil, which is subsequently converted into a thymine through DNA replication or repair (**Fig. 1B-C**). ScBE3-mediated blocking of repair further enhances the editing efficiency (36). Gene disruption can be achieved by converting a cytosine into a thymine to mutate a start codon or to generate a premature stop codon (**Fig. 1D**) (27). We employed CRISPR-Cas9 arrays derived from *Streptococcus pyogenes* (SpCas9) (19, 20), which enable expression of multiple guide RNAs (gRNAs) from one promoter in a single construct, permitting programmable and simultaneous editing of three target genes.

### Temperature-sensitive CRISPR array plasmids allow for sequential rounds of multiplexed base editing

To demonstrate this base-editing approach, we designed a triple-spacer array to direct ScBE3 to introduce premature stop codons in the *S.*Tm genes *sifA, sseJ*, and *sseL*. We selected these genes based on prior literature linking these SPI-2 effectors to development of a stable *Salmonella-*containing vacuole decorated with host cholesterol (37, 38), though it remains unclear how parallel disruption of these three genes would affect virulence. We hypothesised that a triple mutant would show a strong reduction in virulence beyond that seen for single mutants, and that such a strain would prove an effective pilot candidate for testing our base editing strategy. Separately, we designed two more triple-spacer arrays targeting another six SPI-2 effector genes (*sifB, srfJ, srgE, pipB, cigR,* and *steB*), the functions of which remain poorly characterized (6). Co-transformation of *S.*Tm with two constructs encoding a CRISPR array and the ScBE3 editor (**Fig. 1E**) was followed by induction with IPTG and arabinose to induce base-editing, with plasmids maintained by antibiotic pressure (**Fig. 1F**). Colonies were screened by PCR and Sanger sequencing to confirm the desired base-edits. Temperature-sensitive CRISPR array constructs were cleared by incubation at higher temperatures, and plasmids were cleared by culture without antibiotic pressure. This strategy was used to generate a triple mutant *sifA-*Q85X, *sseJ*-Q82X, *sseL*-Q18X (**Fig. 1G**, left), where X denotes a premature stop codon at the indicated amino acid position. Separately, we performed two sequential rounds of editing using two spacer arrays (one targeting *sifB, srfJ,* and *srgE*, and another targeting *pipB, cigR, steB*), resulting in a sixfold mutant *sifB*-Q22X, *srfJ*-Q13X, *srgE*-Q54X, *pipB*-W38X, *cigR*-M1I, *steB*-M1I (**Fig. 1G**, right). Our strategy to sequentially use triple-spacer arrays was based on reports that further upscaling may vary gRNA abundance (39), and speculation that base editor efficiency could be titrated leading to less efficient targeting, similar to what has been proposed for CRISPRi (40). Indeed, during sequencing verification, we observed variation between sampled colonies in which some colonies carried correctly edited codons while others did not (**Fig. 1H**), demonstrating the variable level of editing efficiency between clones. As a final proof of concept, we separately constructed a ninefold mutant by iterative rounds of transformation and editing using four CRISPR spacer arrays targeting twelve different genes in one strain (**Fig. S1**). Interestingly, while nine of twelve intended edits were introduced, we also detected bystander editing activity that resulted in premature stop codons in the target gene *sspH2* which were upstream of the intended edit site. Together, these data show that multiplexed cytidine base editing represents a viable strategy for iterative deletion of a large number of target genes in a single bacterial strain.

### Base-edited and mutant strains show strong attenuation *in vivo*

Next, we investigated the virulence defect of our base-edited triple mutant *sifA-*Q85X, *sseJ*-Q82X, *sseL*-Q18X using an animal model of systemic *Salmonella* infection (**Fig. 2A**) (41). We contrasted the performance of this strain against a mutant strain *S.*Tm Δ*sifA* Δ*sseJ* Δ*sseL*, which we constructed using well-established techniques for recombineering and phage transduction (42–45) and which acquired very few polymorphisms during strain construction (**Table S5**). This approach allows for the comparison of creating mutants by introducing premature stop codons versus deletion of entire genes, and also provides a first means to assess the impact of off-target base edits on bacterial fitness. Mice that were infected with *S.*Tm WT showed very high bacterial loads in the liver (**Fig. 2B**, upper) and spleen (**Fig. 2B**, lower), while a SPI-2 T3SS deficient *S.*Tm Δ*ssaV* mutant failed to replicate in these systemic niches. The base-edited *sifA-*Q85X, *sseJ*-Q82X, *sseL*-Q18X strain was recovered at similar levels than *S.*Tm Δ*ssaV*, while *S.*Tm Δ*sifA* Δ*sseJ* Δ*sseL* was recovered at slightly but significantly higher levels than the base edited strain.

**Figure 2.**
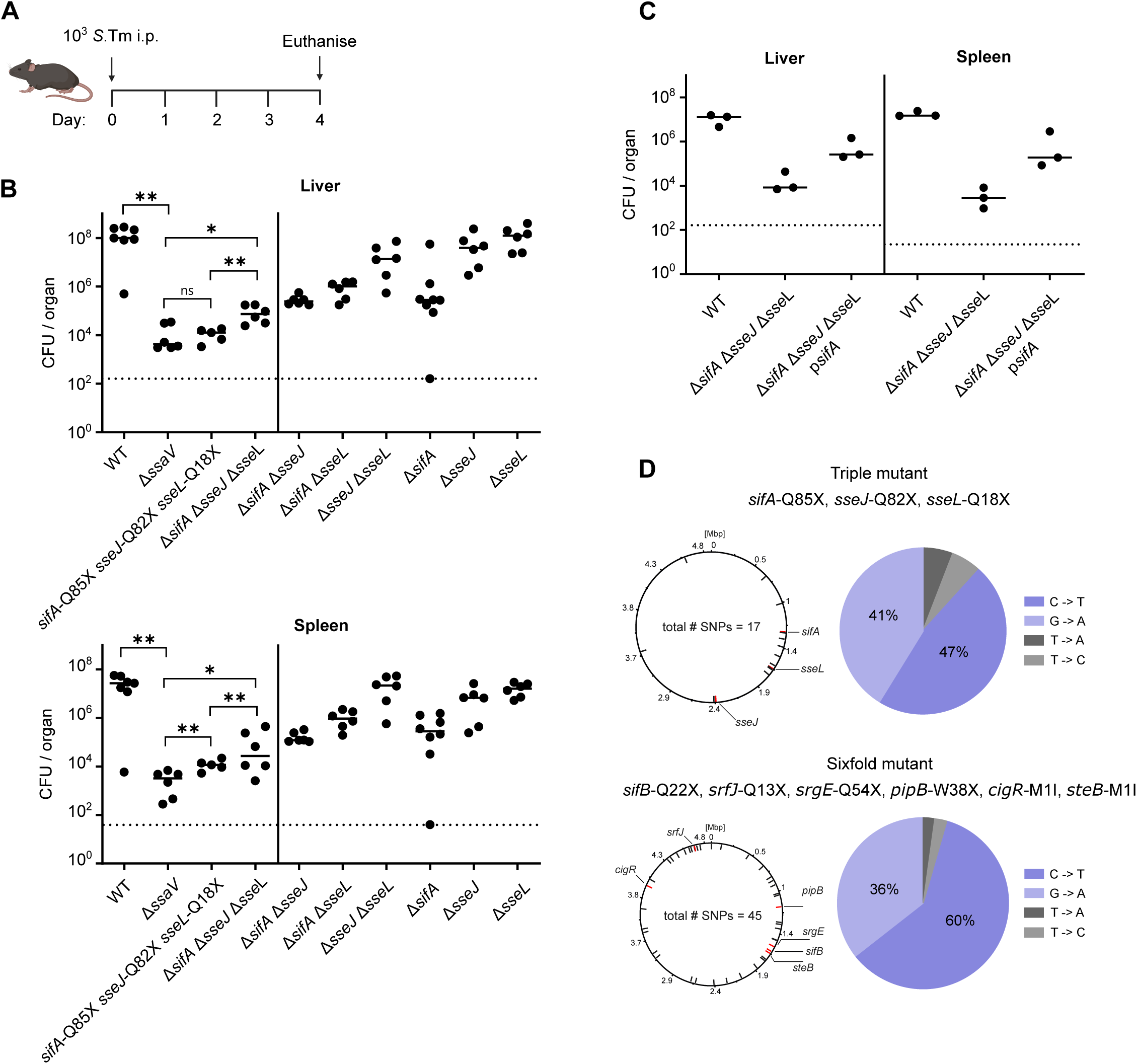
Attenuation of edited and mutant strains in an animal model of infection. **A)** Experimental scheme for systemic infection of mice with *S.* Typhimurium. C57BL/6J Mice were infected by intraperitoneal injection with 10^3^ *S.*Tm. Mice were euthanised at day 4 post infection. Bacterial loads in the liver and spleen were quantified by CFU plating to selective media. **B)** *S.*Tm recovered from the liver (upper) and spleen (lower). Mice infected with strains indicated on x-axis (*n* = 5-8 mice per group). **C)** *S.*Tm recovered from the liver (left) and spleen (right). Mice infected with strains indicated on x-axis (*n* = 3 mice per group). Horizontal bars denote median, dotted lines denote limit of detection. Statistical differences between WT and indicated groups determined by two-tailed Mann Whitney-U test, (p≥0.05 not significant (ns), p<0.05 (*), p<0.01 (**). **D)** Mapped single-nucleotide polymorphisms (SNPs) based on whole-genome sequencing (WGS) of base-edited strains. For both the triple mutant (upper) and sixfold mutant (lower), circular maps show genomic distribution and total number of SNPs detected in black, while intended base-edits are shown in red. Pie charts show percentage of each detected off-target SNP, with putative cytidine deaminase off-target edits in blue and other polymorphisms in grey.

Further, we deployed a series of single and double mutants constructed using classical recombineering to determine which of these three effectors contributed to this strong virulence defect. We observed a trend in which deletion of *sifA* was sufficient to reduce bacterial loads relative to the WT, but any combination of double mutant did not seem to reduce levels to that of the triple mutant (**Fig. 2B**). In support of the keystone role of SifA (46–48), we complemented the *S.*Tm Δ*sifA* Δ*sseJ* Δ*sseL* strain with a plasmid-encoded copy of *sifA*, and observed a partial but not complete rescue of bacterial virulence relative to WT levels (**Fig. 2C**). These data suggest that while SifA plays a cardinal role in bacterial virulence, the deletion of these three effectors together produces a particularly attenuated strain that is impaired almost to the degree of a SPI-2 T3SS deficient strain.

Our observation that a base-edited strain was less virulent then a recombineered mutant may suggest that off-target edits can impact bacterial fitness. Various reports have shown that base editors can also result in undesired edits, either through guide-dependent off-target activity (49), or through high cytidine deaminase activity on single-stranded DNA which is independent of the guide (50–52). However, it remains unexplored whether these mutations can impact bacterial fitness during infection. To assess the degree of off-target edits in our strains, we performed whole-genome sequencing on both the triple and sixfold mutants. In addition to the on-target edits, we observed a range of off-target edits for the triple mutant (*n* = 17, 47% C to T, 41% G to A) and for the sixfold mutant (*n* = 45, 60% C to T, 36% G to A) (**Fig. 2D**, **Table S6**). Given the high proportion of C-to-T variants (or G-to-A for the opposite DNA strand), it seems likely that these mutations are attributable to cytidine deaminase off-target activity, as seen previously when cytidine base editors were used in *Corynebacterium* (52) and *E.coli* (53). These data suggest that cytidine deaminase base editors guided by CRISPR arrays can precisely and efficiently convert target bases but concomitantly introduced off-target changes elsewhere in the genome, in a manner that is faster and simpler than classical recombineering but can result in confounding mutations that impact fitness.

### Cytidine deaminase base editing causes off-target changes across the bacterial genome

Thus far, we observed that CRISPR array-guided cytidine deaminase base editors could precisely convert target bases (**Fig. 1G**) but also introduced a range of off-target base conversions (**Fig. 2D**), and that a base-edited triple mutant was slightly less virulent than a strain in which these same target genes were fully deleted (**Fig. 2B**). These data led us to explore the degree of off-target activity when this editing process is performed in parallel, and how these mutations might impact virulence phenotypes. We deployed a series of isogenic *S.*Tm wild-type strains that each bear a unique genomic tag that can be amplified by RT-qPCR. In parallel, each strain was co-transformed with plasmids encoding the ScBE3 base editor and a CRISPR spacer array designed to target *sifA, sseJ*, and *sseL* (**Fig. 3A**). After one round of base editing, screening, and clearing (as in **Fig. 1G**), this resulted in a set of seven strains each with the desired edits (*sifA-*Q85X, *sseJ*-Q82X, *sseL*-Q18X), and with a unique genetic tag compatible with qPCR amplification. To assess the degree of off-target changes, we performed whole-genome sequencing of these seven strains and analysed polymorphisms relative to the WT parent strain (**Fig. 3B**). While all edited strains showed the correct base conversions in target genes, we also observed a varying degree of off-target changes (from *n* = 28 to *n =* 124) (**Table S7**). As previously (**Fig. 2D**), the majority of detected SNPs (>92%) were C-to-T or G-to-A edits, suggesting these arose due to cytidine deaminase off-targeting. Given that editing conditions and CRISPR array design were the same for all strains, these data suggest an inherent degree of variability in the extent of off-target enzyme activity under these conditions.

**Figure 3.**
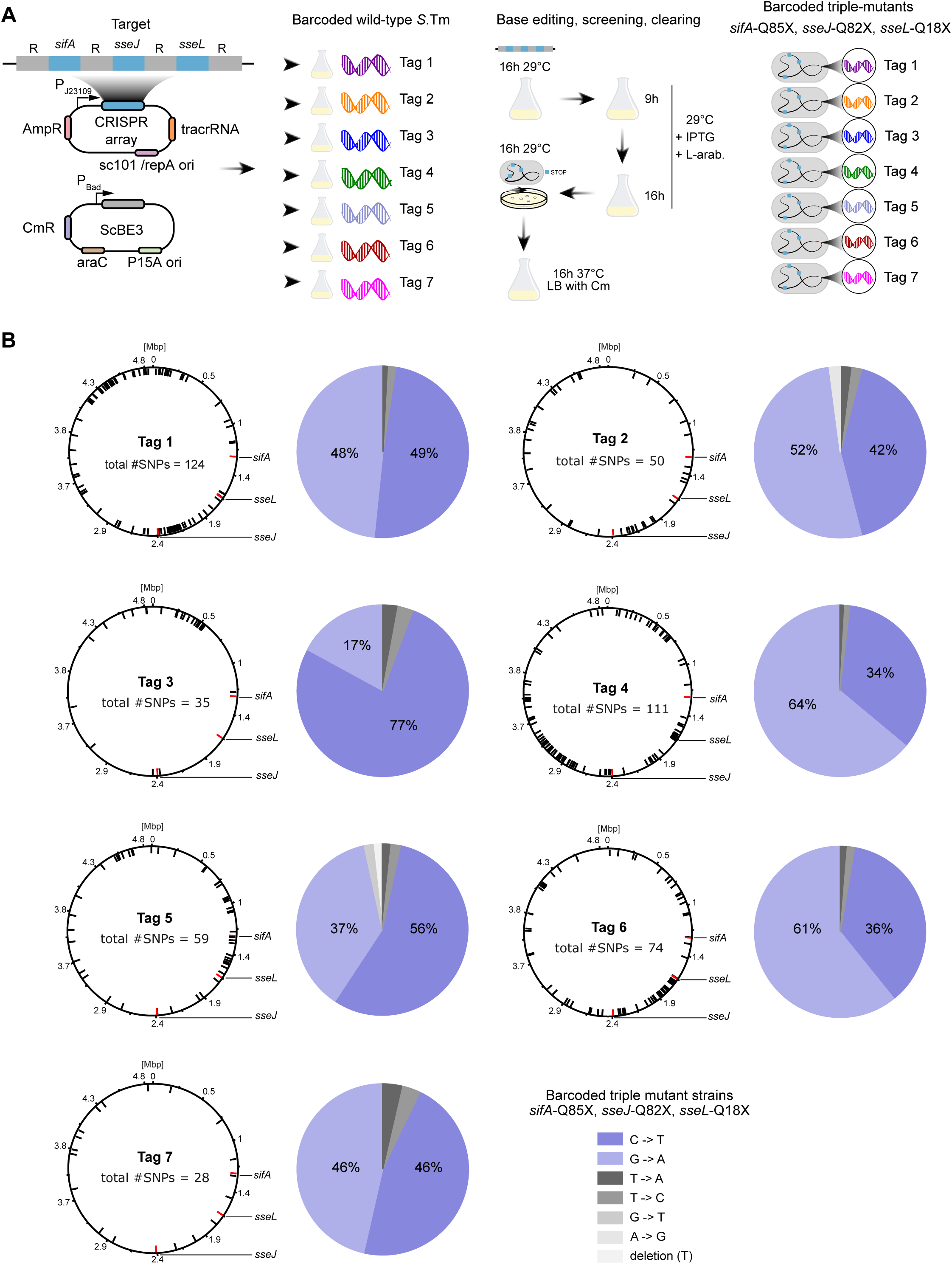
Parallel base-editing of isogenic tagged *S.*Tm strains results in variation of off-target changes. **A)** Experimental overview of procedure for parallel editing of tagged strains. Seven isogenic wild-type *S.*Tm strains bearing unique tags inserted in a fitness neutral position in the genome were co-transformed with plasmids bearing the CRISPR array and base editor. Base editing, screening, and clearing of constructs was performed as previously (Fig. 1). Resulting strains were verified by Sanger sequencing to confirm the desired base edits to target genes, and to confirm no disruption to genomic tags. **B)** Mapped single-nucleotide polymorphisms (SNPs) based on whole-genome sequencing (WGS) of seven base-edited genome-tagged strains. Circular maps show genomic distribution and total number of SNPs detected in black, while intended base-edits are shown in red. Pie charts show percentage of each detected off-target SNP, with putative cytidine deaminase off-target edits in blue and other polymorphisms in grey.

### Off-target edits have a variable impact on bacterial virulence *in vivo*

We next examined how the phenotype of our seven base-edited tagged strains might be affected by off-target changes to the genome. During *in vitro* growth assays, we observed no apparent difference in growth kinetics compared to the WT parent strain (**Fig. 4A**). Similarly, co-culture of all seven tagged strains followed by genomic DNA extraction and qPCR analysis showed a relatively similar proportion of each tag within the total population (**Fig. 4B**). These data suggested that bacterial fitness *in vitro* was not affected as a result of base editing. Next, we assessed the virulence of these strains *in vivo* by infecting mice with a mixed inoculum comprising the seven tagged and edited strains, alongside tagged versions of *S.*Tm WT and *S.*Tm Δ*ssaV* (**Fig. 4C**). This infection model permits highly sensitive measurements of the relative fitness of a strain within the experimental context of one animal. Here, we reproducibly observed dominance of *S.*Tm WT within the spleen of infected animals, while *S.*Tm Δ*ssaV* was severely outcompeted, as expected. For the base-edited strains, we observed a spectrum of phenotypes in which some edited strains were out-competed to the degree of *S.*Tm Δ*ssaV* (particularly strains bearing Tag 1 and Tag 5), while the other strains had less pronounced defects (**Fig. 4D, Fig. S2A**). These data may suggest that variation in which genes are affected by off-target cytidine deaminase activity may produce variation in virulence phenotypes.

**Figure 4.**
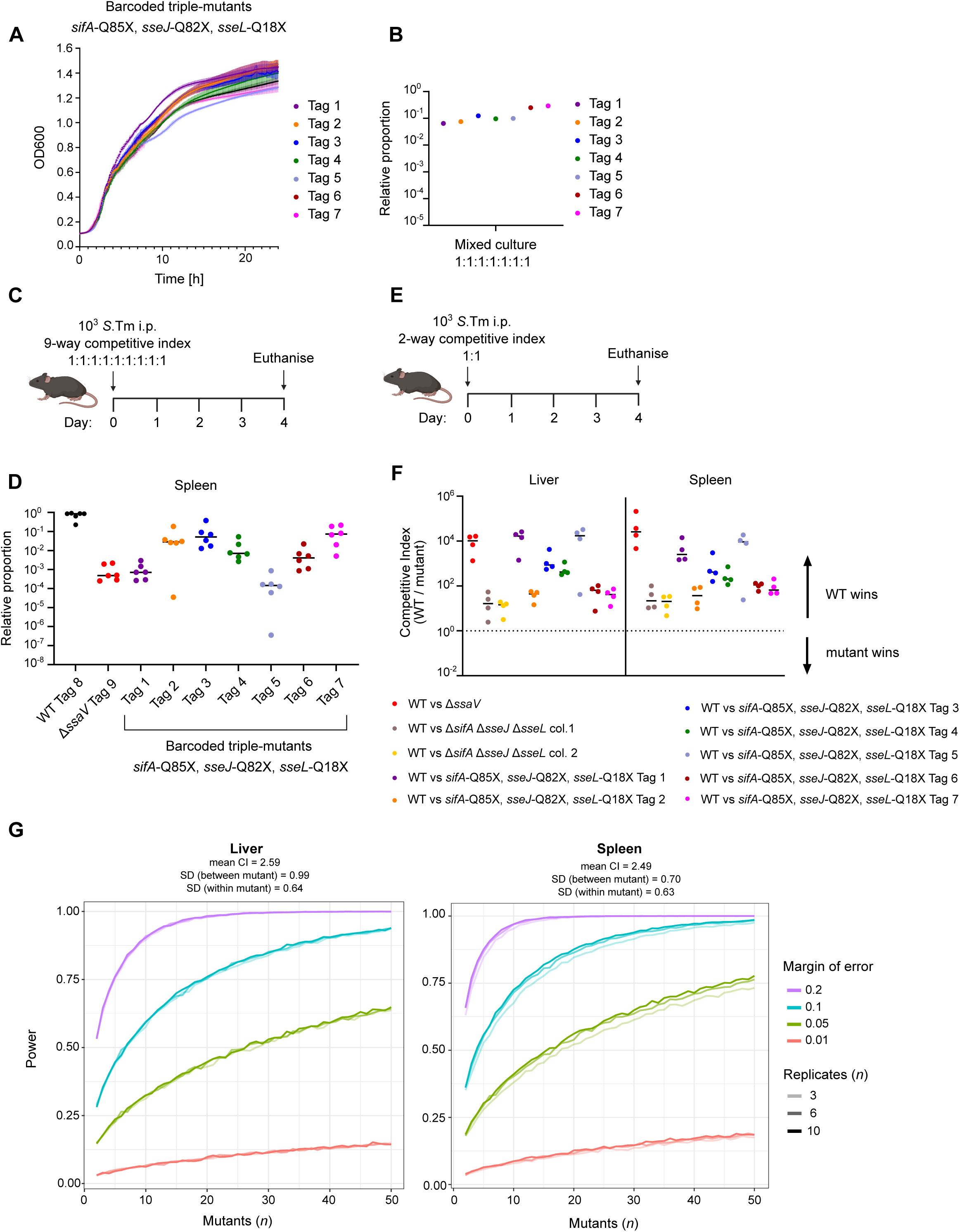
Variation in clonal fitness arising from off-target edits can be surmounted by sufficiently-powered multi-clone testing. **A)** Growth curves for barcoded base-edited triple mutants shows no difference in growth when strains are cultured in minimal medium for 24 hours. **B)** Relative proportion of each tagged strain in a mixed culture after 12 hours incubation in LB media. Proportions determined by qPCR analysis of tag abundance following DNA extraction from liquid culture. **C)** Experimental scheme for 9-way competitive index infection of mice. Mice were injected with an infectious dose of 10^3^ *S.*Tm comprising equivalent volumes of nine strains (shown in Fig. 4D). Mice were euthanised at day 4 post infection, and spleen samples were homogenised and used to inoculate overnight enrichment cultures. **D)** Relative proportion of each tagged strain within the total population, calculated by qPCR analysis of samples from Fig. 4C. Proportions are calculated for populations from one animal, then sorted by bacterial strain (x-axis). (*n* = 6 mice). **E)** Experimental scheme for 2-way competitive index infection of mice. Overnight cultures of strains (indicated in Fig. 4F) were mixed 1:1 and diluted to achieve an infectious dose of 10^3^ *S.*Tm, which was delivered to mice by intraperitoneal injection. Mice were euthanised at day 4 post infection. **F)** Competitive index of indicated strains determined by CFU plating to selective media (Cm for WT, Amp for mutant strains). Competitive index calculated by dividing CFU for WT by CFU for mutants. Indices calculated for bacteria recovered from the liver (left) and spleen (right) of mice. Dotted line denotes a neutral competitive index *i.e.* no difference between two indicated strains. **G**) Estimation of the number of mutants constructed by base editing (x-axis) needed to achieve indicated levels of power with allowable margins of error. Simulations derived from mutant fitness CFU data shown in Fig. 4F and performed for values derived from liver (left) and spleen (right). Power, estimated from a simulation-based approach, was defined as the proportion of simulations where the mean fitness of the simulated dataset fell within a specified margin of error (MOE) of the observed mean fitness. Fitness data was simulated under a linear mixed effects model with mutant identity as the random effect, with mean and variances taken from the log10-transformed *in vivo* competition index data. 10000 simulations for each combination of number of mutant clones and replicates per clone were performed. For liver simulations: mean CI = 2.59, SD (between mutant) = 0.99; SD (within-mutant) = 0.64. For spleen simulations, mean CI = 2.49; SD (between mutant) = 0.70; SD (within-mutant) = 0.63.

Next, we performed a series of one-to-one competitive index infections, in which each edited strain was competed against *S.*Tm WT (**Fig. 4E**). Separately, we assessed two clones of *S.*Tm Δ*sifA* Δ*sseJ* Δ*sseL* (constructed by traditional recombineering) in order to establish the baseline competitive index for this triple mutant. As expected, *S.*Tm Δ*ssaV* was severely outcompeted by *S.*Tm WT in both the liver and spleen, while both clones of *S.*Tm Δ*sifA* Δ*sseJ* Δ*sseL* were more modestly outcompeted (**Fig. 4F, Fig. S2B**). Consistent with our data from a 9-way co-infection (**Fig. 4D**), we found that both edited strains bearing Tag 1 and Tag 5 were outcompeted by *S.*Tm WT, to a degree similar to that seen for *S.*Tm Δ*ssaV* (**Fig. 4F, Fig. S2B**). Other tagged strains (Tags 2, 6, and 7) performed in a manner that was strikingly similar to the recombineered clones of *S.*Tm Δ*sifA* Δ*sseJ* Δ*sseL*, while some tagged strains showed an intermediate phenotype (Tags 3 and 4). These data suggest that while base-editing can produce strains that phenocopy a recombineered mutant, the spectrum of off-target effects introduced during base editing can further decrease bacterial virulence, even to the level of a mutant fully defective for intracellular infection.

### Predicting sample sizes for multi-clone testing that may permit accurate phenotype discovery

Finally, we explored how we can employ a series of edited clones carrying diverse sets of background mutations to determine the phenotypes associated with the mutations of interest. Our *in vivo* data (**Fig. 4F**) indicated that several edited strains (Tags 2, 6, and 7), showed similar levels of attenuation to the recombineered clones (*S.*Tm Δ*sifA* Δ*sseJ* Δ*sseL*; col. 1, col. 2). Thus, parallel editing produces some clones that will show the correct phenotype, while other edited clones are more strongly attenuated by their respective background mutations. A viable strategy for correctly deducing an unknown phenotype may be to analyse a sufficient number of edited clones, each of which carry a different set of background mutations. To explore this concept, we performed a statistical power analysis to estimate the number of edited mutants needed to accurately determine mutant fitness. Competitive index data from our *in vivo* experiments (**Fig. 4F)** was used to estimate the variability in fitness across barcoded mutant clones (see **Methods**). We simulated fitness data (**Fig. 4G**) using these estimates for different numbers of hypothetical edited clones (*n =* 2-50) and for different numbers of experimental replicates per individual edited clone (*n =* 3-10). For each of these simulated datasets, we estimated the fitness of hypothetical mutants. This analysis revealed that fitness estimation accuracy depended primarily on the number of mutant clones rather than the number of replicates per clone. Increasing the number of clones markedly improved power, particularly at higher allowable margins of error (ME). For example, in the liver data at a strict ME = 0.01, power remained low (0.14) even with 50 clones and 10 replicates, whereas at a more permissive ME = 0.2, power reached 1.0 under the same conditions (**Fig. 4G**, left). Power gains started to slow down beyond a certain number of mutant clones, which can be seen clearly at ME = 0.2, where beyond approximately 24 clones, there is no further improvement in power. Similar patterns can be observed in the spleen data, where fewer mutant clones are required to achieve similar power due to the lower between-mutant variance (**Fig. 4G**, right). Together, these simulations suggest that even with a permissive margin of error, a large number (*n >* 10) of base-edited clones would be needed to accurately determine unknown phenotypes, while at a strict margin of error an even greater number of clones are necessary (*n* > 50). While testing this number of independent clones is not a viable strategy in some experimental contexts, it is possible to simultaneously interrogate dozens of clones by exploiting DNA tagging coupled with high-throughput population analyses such as RT-qPCR or sequence counting (54–57).

Ultimately, our data show that cytidine deaminase base editors can efficiently produce complex multi-mutant strains, but that off-target activity results in genetic changes that may confound phenotypic characterisation. This drawback can be overcome by testing a sufficient number of independently constructed clones of a multi-mutant strain in high-throughput experimental approaches.

## Discussion

Understanding how genetic interdependencies shape phenotypic outcomes requires tools for genetic engineering that are robust and efficient. Genetic manipulation of many clinically relevant pathogens, including *Salmonella enterica, Listeria monocytogenes,* and *Staphylococcus aureus,* has led to a deeper understanding of how these bacteria undermine host processes to achieve infection (5, 58, 59), but technology to engineer bacteria remains relatively low-throughput and is characterised by a range of disadvantages. Here, we explored the use of CRISPR array-guided cytidine deaminase base editors as a programmable, scalable, and efficient means of disrupting multiple bacterial genes. We used this approach to understand the contribution of three SPI-2 T3SS genes to systemic *S.*Tm infection in mice, and demonstrated a framework for assessing complex virulence phenotypes against a background of off-target cytidine deaminase-mediated mutations that confound phenotypic characterisation.

A particular advantage of this approach is the incorporation of three spacers into a plasmid-encoded CRISPR array, which permits the simultaneous disruption of three target genes (**Fig. 1**). Further, by employing a temperature-sensitive version of the multi-spacer CRISPR array plasmid and performing multiple rounds of transformation, editing, and clearing, it is possible to rapidly scale the number of disrupted effector genes. In contrast, the well-established method of engineering *Salmonella* genomes via lambda red recombineering and P22 transduction is limited to one gene (or one genetic region comprising chromosomally co-located genes), relies on incorporation of resistance cassettes which must first be PCR-amplified to be homologous to the target gene, and results in genetic scars in the chromosome after removal of resistance cassettes by FLP recombinase excision (42). Thus, gene disruption by multiplexed base editing offers a more efficient means of disrupting multiple genes simultaneously, which we demonstrate here by generation of triple-edited strain (*sifA*-Q85X, *sseJ*-Q82X, *sseL*-Q18X) and a sixfold-edited strain (*sifB*-Q22X, *srfJ*-Q13X, *srgE*-Q54X, *pipB*-W38X, *cigR*-M1I, *steB*-M1I) (**Fig. 1G**). As the plasmids encoding both the CRISPR array and base editor are easily cleared by temperature shifts and withdrawal of antibiotic pressure, we anticipate that further rounds of editing could be used to disrupt even more target genes within any given strain, as we achieve with a ninefold-edited strain (**Fig. S1**). Finally, as the editing reaction contains many independently edited clones, we speculate that generating and experimentally testing many clones of a multi-edited strain may represent an alternative means of experimentally proving a genetic-phenotypic relationship. This may provide a useful contrast to the classical approach of complementing a gene deletion in one strain to satisfy the molecular Koch’s postulates (60), which is logistically difficult when multiple genes are deleted in one strain.

However, various drawbacks are associated with cytidine deaminase base editors. We observed between-colony variation in editing efficiency, such that several colonies needed to be verified by PCR and sequencing to achieve a correctly edited triple mutant, while screening of up to 10 colonies was required for the sixfold mutant to isolate a strain with all six desired edits (**Fig. 1G-H**). These data support our strategy for using CRISPR arrays targeting only three genes in one round of editing, especially given prior reports describing variation in editing efficiencies due to the identity of individual guides comprising the CRISPR array (39). We also report a significant degree of mutations that likely arose from off-target cytidine deaminase activity (**Fig. 2D**, **Fig. 3B**). Base editors of this class have been shown to lead to Cas9-independent off-target mutations in mammalian cells (50), plants (51), and bacteria (52) through random cytidine deaminase activity on single-stranded DNA (61). Here, we explored the degree of off-target activity by parallel editing of seven isogenic wild-type *S*.Tm strains (**Fig. 3B**), and observed large variation in the number of mutations arising (*n* = 28 to 124 SNPs per strain). These edited strains showed similar growth kinetics *in vitro* (**Fig. 4A-B**) but produced a variation of virulence phenotypes in animal models of infection (**Fig. 4D**, **Fig. 4F**), likely due to the random nature of off-target activity disrupting genes that seem to be critical for virulence (**Table S7**). While some edited strains produce a virulence defect similar to classically-recombineered mutant strains (*n* = 3/7) (**Fig. 4F**), our simulated data suggests that many clones of edited strains would be needed to accurately characterise unknown phenotypes of target genes (**Fig. 4G**). Nonetheless, a single base-editing reaction culture theoretically contains a great number of correctly edited clones, and the application of whole-genome sequencing to carefully select for minimal off-target effects could prove a useful strategy. Importantly, cytidine base editors have gained traction as a technology for genetic engineering (52, 53) and for genome-wide screens (62–64), and here we describe how the scope of off-target activity can variably impact strain fitness during infection, which might confound these experimental approaches.

The reduction of off-target mutations is a critical aspect that lies ahead of the broader application of cytidine base editing technologies. Modulation of high cytidine deaminase activity represents a viable strategy, and could be achieved through the use of virally-derived anti-deaminase proteins (65) , by the use of cytidine base editors with mutated deaminase domains (53, 66), or by fusions with a cleavable deaminase inhibitor (67). Further, new variants of cytidine base editors based on an engineered TadA domain instead of a cytidine deaminase domain have been shown to enable robust target editing while avoiding detectable off-target mutations in mammalian cells (68). Other strategies explore mitigation of Cas9-dependent off-target mutations, through use of anti-CRISPR proteins that interfere with target DNA binding or cleavage, crRNA loading, or effector complex formation (69–72). However, many of these strategies have been applied for reducing off-target editing in mammalian cells, and it remains to be shown whether efficiencies in editing bacterial genomes can be similarly improved. Incorporation of one or more of these elements may prove useful in minimising the number of off-target effects, which could be tested using the parallel editing and sequencing strategy deployed here.

As a proof of concept, we elected to introduce premature codons into three genes of *Salmonella* Typhimurium encoding effectors of the SPI-2 T3SS. A body of literature supports the key role played by SifA in stabilising and expanding the *Salmonella*-containing vacuole (SCV) (46–48, 73), while recent reports suggest a cooperative role for SseJ and SseL in orchestrating the deposition of host cholesterol onto the SCV membrane (37, 38). Here, we describe a strong virulence defect for a *sifA/sseJ/sseL* mutant strain, both when constructed by recombineering and by base editing. Our data suggests that abrogation of these three genes alone reduces a mutant strain close to the level of a SPI-2 T3SS deficient *S.*Tm Δ*ssaV* mutant (**Fig. 2B**). Further work is needed to characterise the interplay between these three effectors, particularly regarding the stability and decoration of the SCV, which reportedly is either destabilised or re-stabilised in mutant strains deficient for various combinations of SPI-2 effectors (37, 47, 74, 75). Mutants which cannot stabilise or expand the SCV are highly impaired for intracellular replication and survival (48), and further characterisation of how these effectors functionally cooperate is warranted.

Finally, we speculate on the study of individual genes with respect to the broader context of genomic integrity. While selective pressures and replication fidelity typically preserve genetic information, a critical aspect of evolutionary adaption is a certain degree of tolerance for genetic changes. The study of host-pathogen interactions historically calls for the characterization of phenotypic traits by the clean deletion and subsequent complementation of a gene in an otherwise constant genetic background. It is interesting to reflect that in nature, the function of a gene occurs in the broader context of genetic plasticity elsewhere in the genome (via selective pressures acting on infidelity in genome replication or various mechanisms of horizontal gene transfer). The use of base editor enzymes to broadly and randomly introduce genetic changes into a bacterial chromosome may prove useful in exploring how genes and their proteins maintain function despite a changing genetic environment. Further, it may be possible to uncover novel genetic interdependencies between genes that have not been detected by techniques that prioritise genomic fidelity. As very high numbers of bacterial cells can be simultaneously edited in one base editing reaction culture, it may be possible to exploit the off-target nature of cytidine deaminase editors for generation of a large number of diversely edited genomes, which may prove useful in the study of genomic plasticity.

Ultimately, we describe how CRISPR array-guided cytidine deaminase base editors provide a means of editing multiple target bacterial genes in a manner that is scalable, programmable, and efficient. The impact of off-target activity on bacterial fitness should more broadly be considered when using base editors of this class. The range of off-target effects we describe calls for routine use of whole-genome sequencing, which will also be helpful in the development of refined or optimised base editor technologies. Such approaches will be useful for the robust and simultaneous disruption of many target genes, a strategy that is necessary to more broadly understand how genetic co-dependencies shape outcomes in host-pathogen interactions.

## Methods

### Ethics statement

C57BL/6 mice originated from breeders originally obtained from Jackson laboratories (catalog no. JAX:00066). Mice were specific pathogen-free (SPF) and bred under full barrier conditions in individually ventilated cage systems in the EPIC mouse facility of ETH Zürich, Switzerland. Mice were held under SPF conditions at the EPIC facility at ETH Zürich (light:dark cycle 12:12 h, room temperature 21 ± 1°C, humidity 50 ± 10%). All animal experiments were reviewed and approved by Tierversuchskommission, Kantonales Veterinäramt Zürich under license ZH158/2019 or ZH108/2022, complying with the cantonal and Swiss legislation. Experiments were performed with 8- to 12-week-old mice of both sexes, and mice were randomly assigned to experimental groups.

### Strains, plasmids and growth conditions

All strains, plasmids and primers are listed in supplementary **Tables S1-3**. Primers were synthesized by Integrated DNA Technologies or Microsynth. Plasmids were propagated and maintained in *E. coli* TOP10 and experiments were carried out using *S.* Typhimurium SL1344 or its genomically barcoded derivatives. *E. coli* and *S.* Typhimurium cells were grown in Luria Bertani (LB) medium (10 g/l NaCl, 5 g/l yeast extract, 10 g/l tryptone) at 37 °C, or 29 °C for strains harboring temperature-sensitive constructs, with shaking at 250 rpm. The antibiotics ampicillin, kanamycin, and/or chloramphenicol were added at 50 µg/ml, 34 µg/ml, and 50 µg/ml, respectively to maintain the plasmids.

### Construction of clean deletion and barcoded *S*.Tm strains

*S.*Tm deletion and barcode-tagged mutants were constructed via lambda red recombination as described in (42) and P22 transduced (76) into the clean SB300 background. The unique 40 bp barcode-tags with the antibiotic cassette (*tag::amp* for WISH tags, *tag::cm* for WITS-1 tag, *tag::kan* for WITS-2 tag) were integrated between the pseudogenes *malX* and *malY* on the SL1344 chromosome (FQ312003.1: at base 1,495,714) (77). The sequences of the seven WISH tags and the WITS-1 and WITS-2 tags are listed in supplementary **Table S4**.

### Cloning of CRISPR arrays

The CRISPR arrays were cloned according to the CRATES method (39). Briefly, forward and reverse oligonucleotides encoding one repeat, one spacer and a 4-nt junction were p5′ phosphorylated with polynucleotide kinase (New England Biolabs) and annealed to form a dsDNA fragment with 4-nt overhangs at the 5′ and 3′ terminal end. For the Golden Gate reaction, 2 µl T4 ligation buffer (New England Biolabs) , 1 µl of each inserts, 50 ng of backbone plasmid, 1 µl of T4 ligase (New England Biolabs) and 1 µl of BsmBI-HFv2 (New England Biolabs) were mixed with the appropriate amount of water to a final volume of 20 µl. A thermocycler was used to perform 25 cycles of digestion and ligation (42 °C for 2 min, 16 °C for 5 min) followed by a final digestion step (60 °C for 10 min), and a heat inactivation step (80 °C for 10 min). The ligation mix (2.5 µl) was used to transform chemically competent *E. coli* TOP10 cells (25 µl). After recovery in SOC medium for 1 h at either 37 °C, or 29 °C for strains harboring temperature-sensitive constructs, with shaking at 250 rpm, cells were plated on LB agar plates with the appropriate antibiotics and incubated overnight at 37 °C or 29 °C. White colonies were chosen for screening and verified by Sanger-sequencing of the PCR-amplified array region.

### Base-editing to disrupt *S*.Tm effector genes

Electrocompetent SL1344 cells were co-transformed with the base-editor and the CRISPR-array encoding plasmids and recovered in SOC medium for 1 h at either 37 °C, or 29 °C for strains harboring temperature-sensitive constructs, with shaking at 250 rpm. After recovery, cells were plated on LB agar plates with the appropriate antibiotics and incubated overnight at 37 °C or 29 °C. A single colony was selected for the base-editing process and used to inoculate LB medium with the appropriate antibiotics and incubated overnight at 37 °C or 29 °C. The next morning the culture was back-diluted 1:1000 in fresh medium with 1 mM IPTG and 20 mM L-arabinose and incubated at 37 °C or 29 °C, with shaking at 250 rpm for 9 h. To increase the editing efficiency this step was repeated for an incubation period of another 16 h (optional to add further 16-h incubation periods). The culture was then plated on LB agar plates with appropriate antibiotics, incubated overnight at 37 °C or 29 °C and obtained colonies were analyzed by colony PCR and Sanger-sequencing of the PCR-amplified genomic locus with the base-editor target site.

For multiple rounds of editing, temperature-sensitive variants of the CRISPR array encoding plasmid were cleared from strains by culturing in LB medium with ampicillin and chloramphenicol but without kanamycin (for editing of WISH-tagged strains depicted in **Fig. 3**), or by culturing in LB medium with chloramphenicol and either ampicillin or kanamycin, as required (for editing of strains depicted in **Fig. 1** and **Fig. S1**). Cultures were incubated overnight at 37 °C with shaking at 250 rpm. The next day, the culture was plated on LB plates with ampicillin and chloramphenicol or ampicillin, chloramphenicol and kanamycin and incubated at 37 °C for 16 h to confirm loss of the temperature-sensitive plasmid. The resulting colonies were made electrocompetent and transformed with the next temperature-sensitive CRISPR-array encoding plasmid targeting a new set of effector genes. With regard to the nomenclature of base-edited genes, for instance, the conversion of the triplet encoding glutamine (Q) at position 85 in the *sifA* gene into a stop codon (X) is designated as *sifA*-Q85X. Similarly, a disruption of a start codon (methionine = M), for example, in the *cigR* gene is designated as *cigR*-M1I.

### Growth experiments

To evaluate the growth of the *S.*Tm mutants (**Fig. 4A)**, four individual single clones per strain were used to inoculate 5 ml LB medium and cultured for 16 h at 37°C, while shaking at 250 rpm. The next morning, the overnight cultures were used to inoculate 200 µl of LB medium supplemented with the appropriate antibiotics on a 96-well plate to a final OD_600_ = 0.01. The plates were incubated in a BioTek Synergy H1 plate reader at 37 °C with shaking, reading the OD_600_ every 3 min for 24 h.

### Mouse infection

Animal infections were performed using genetically susceptible C57BL/6J mice. Bacterial cultures were incubated for 12 h at 37 °C on a rotating wheel, then washed several times in cold PBS to remove culture media and antibiotics. Cultures were diluted to achieve an infectious inoculum of approximately 1000 CFU in 100 µl of PBS. Mice were injected intraperitoneally and monitored for adverse effects following injection. Feces were collected at the indicated time points. Mice were sacrificed and organs were harvested at the end of the infection. For plating, the samples were homogenized with a steel ball in a tissue lyser (Qiagen) for 2 minutes at 25 Hz frequency. The homogenized samples were diluted in PBS, plated on MacConkey (Oxoid) plates supplemented with the relevant antibiotics, and incubated at 37 °C overnight. Colonies were counted the next day and represented as CFU per organ, as required.

The competitive index (CI), described in **Fig. 4F**, calculates the change in the ratio of two bacterial strains (A and B) from the beginning of the infection experiment to the end, considering their relative abundance within the host. A CI value greater than one indicates that strain A has a competitive advantage over strain B in the host environment, while a CI value less than 1 indicates the opposite.

### Whole genome sequencing and variant calling

Genomic DNA was extracted from overnight enrichment cultures inoculated with samples of organs or feces from infected animals. Genomic DNA was extracted from 1 ml of overnight culture using a QIAamp DNA Mini Kit (Qiagen). Illumina MiSeq sequencing operated by Novogene (Cambridge) was performed to generate 150 bp paired end reads with at least 50× coverage across the genome.

To analyze single nucleotide variants (SNVs) in the base-edited strains, we used the Geneious Prime® 2023.2.1 software. The Geneious mapper was used with medium-low sensitivity and three iterations to align the FASTQ files of paired-end reads obtained from WGS against the reference sequence of *S.*Tm SL1344 (FQ312003.1). Variants were called with a minimum coverage of 10 and a sensitivity of 70%, resulting in variant frequencies of 90-100% for most SNVs detected (**Table S5-6**). Default settings were used for all other parameters. The analyzed strain CBS-1378 still carried the base-editor construct encoding the *araC* gene, which is also encoded on the SL1344 genome. To avoid erroneous SNV calls, the araC gene was excluded from the analysis. The chromosomal distribution of SNVs was depicted on circular plots generated with DNAPlotter (78).

### Quantification of barcoded (WISH-tagged) *Salmonella* by qPCR

Genomic DNA was extracted from overnight enrichment cultures inoculated with samples of organs or feces from infected animals. Genomic DNA was extracted from 1 ml of overnight culture using a QIAamp DNA Mini Kit (Qiagen). qPCR analysis was performed as described previously (77, 79). Briefly, tag enumeration was achieved using RT-qPCR using the FastStart Universal SYBR Green Master (Rox) (Roche, 13206900) mixed with primers specific to each WITS or WISH tag. The relative proportion of tags within a sample was calculated by dividing the estimated number of tag copies for a given tag by the sum of tag copies for all tags.

### Power analysis

A simulation-based power analysis was performed to quantify to what extent off-target mutations in the barcoded mutant clones confound the estimation of mutant fitness. The observed mean, between-clone variance, and within-clone (replicate level) variance of mutant fitness was first estimated by fitting an intercept-only linear mixed effects model to the log10-transformed *in vivo* competition assay data, with mutant clone identity as the random effect. These estimates were then used to forward-simulate fitness data for different combinations of mutant clones (2–50) and replicates per clone (2–10) under the same mixed-effects structure to understand how sample size affects power. For each combination, 10000 simulations were run, and the estimated mean fitness from each simulated dataset was compared to the observed mean fitness. Power was defined as the proportion of simulations where the estimated mean fell within a specified margin of error (0.01, 0.05, 0.1, or 0.2) of the observed mean.

### Statistical analyses

All statistical analyses were performed using a two-tailed Mann Whitney-U test. P-value (P) > 0.05 is shown as ns, P < 0.05 is shown as *, P < 0.01 is shown as **.

## Acknowledgements

We thank Scott Collins for providing the base editing plasmid SPC914 and Timothy Duchow for support with Geneious. We thank Prof. Michael Hensel for the gift of p2104. We would also like to thank the staff at RCHCI and EPIC animal facilities for their support.

## Contributions

J.P.M.N., S.G., C.L.B., and W.D.H. conceived the study. J.P.M.N., S.G., S.S., R.R.R., C.L.B., and W.D.H. designed the experiments. J.P.M.N., S.G., and S.S. performed experiments and analysed data. J.P.M.N., S.G., S.S., R.R.R., C.L.B., and W.D.H. contributed to writing and editing the manuscript.

---

**Supplementary Figure 1.**
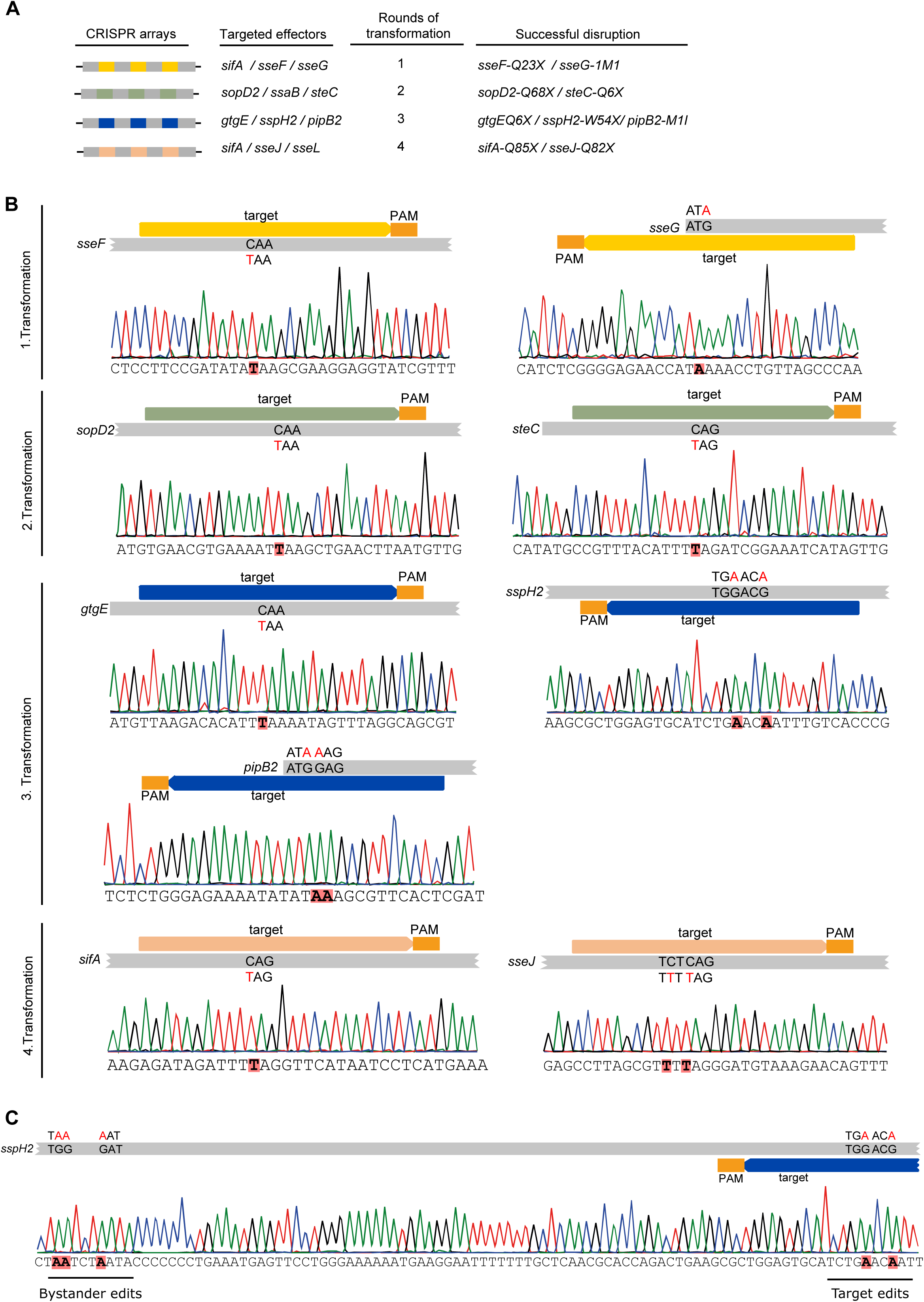
Multiplexed editing of target bases in nine different genes in one *S.*Tm strain via iterative rounds of editing. **A)** Design of CRISPR spacer arrays targeting 12 genes encoding *S.*Tm SPI-2 T3SS effectors. Rounds of transformation, editing, clearance, and sequence verification were performed as in Fig. 1. Four rounds of transformation using different CRISPR arrays resulted in successful disruption of indicated genes at given codon positions. **B)** Verification of edited target bases by colony PCR and Sanger sequencing. Target bases are indicated in upper sections of each panel, while chromatograms below indicate edited bases highlighted in red. **C)** Detection of off-target activity resulting in unintended introduction of premature stop codons in target gene *sspH2* during third round of transformation. Bases in red show intended edits (right side) while bystander edits upstream are also highlighted (left side).

**Supplementary Figure 2.**
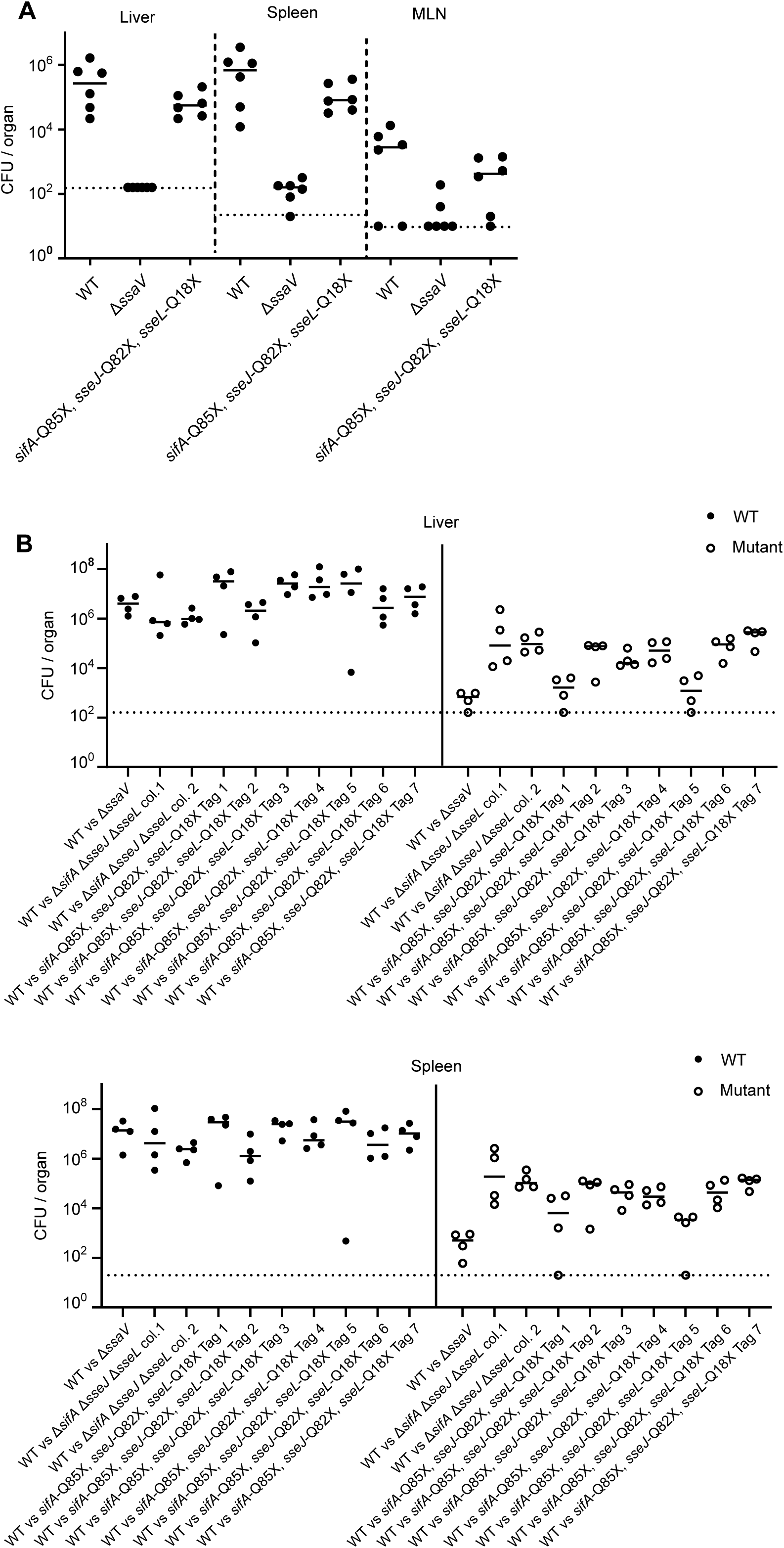
Absolute CFU of *S.*Tm strains used in competitive index infections. **A)** Total CFU of indicated strains from mice infected in 9-way competitive index infections (Fig. 4D). CFU determined by plating to selective media containing: 15 µg/ml chloramphenicol (for WT), 50 µg/ml kanamycin (for Δ*ssaV*); or 50 µg/ml ampicillin (for base-edited mutants). CFU per organ shown for liver (left), spleen (middle), and mesenteric lymph node (MLN, right). **B)** Total CFU of indicated strains from mice infected in 2-way competitive index infections (Fig. 4F). CFU determined by plating to selective media containing 15 µg/ml chloramphenicol (for WT) or 50 µg/ml ampicillin (for indicated mutant strain). CFU per organ shown for liver (upper) and spleen (lower). Dotted lines denote respective limit of detection for each organ, horizontal bars denote median.

**Table S1.** List of microbial strains.

| ID | Strain derivate | Selective marker | Description | Plasmids | Source |
| --- | --- | --- | --- | --- | --- |
| pCB2 | Escherichia coli TOP10 | StrR | mcrAΔ(mrr-hsdRMS-mcrBC) Φ 80lacZ<br>ΔM15 ΔlacX74 deoR<br>recA1araD139Δ(ara-leu)7697 galU galK<br>rpsL endA1 nupG |  | Invitrogen |
| Z1629 | S.Tm SL1344 | SmR | SB300 WT |  | (80) |
| M2730 | S.Tm SL1344 | SmR | ΔssaV |  | (81) |
| Z6695 | S.Tm SL1344 | KanR | ΔsifA ΔsseJ ΔsseL |  | This study |
| Z8235 | S.Tm SL1344 | KanR / CmR | ΔsifA ΔsseJ |  | This study |
| Z6683 | S.Tm SL1344 | SmR | ΔsifA ΔsseL |  | This study |
| Z8237 | S.Tm SL1344 | Kan / CmR | ΔsseJ ΔsseL |  | This study |
| Z5699 | S.Tm SL1344 | KanR | ΔsifA |  | This study |
| Z6531 | S.Tm SL1344 | KanR | ΔsseJ |  | This study |
| Z6546 | S.Tm SL1344 | CmR | ΔsseL |  | This study |
| T2896 | S.Tm SL1344 | AmpR | ΔsifA ΔsseJ ΔsseL with complemented<br>P<sub>sifA</sub> sifA | p2104 | This study |
| T2897 | S.Tm SL1344 | AmpR | ΔsifA ΔsseJ ΔsseL with complemented<br>P<sub>sifA</sub> sifA | p2104 | This study |
| CBS-2871 | S.Tm SL1344 | AmpR | WISH-1 |  | (56) |
| CBS-2872 | S.Tm SL1344 | AmpR | WISH-3 |  | (56) |
| CBS-2873 | S.Tm SL1344 | AmpR | WISH-4 |  | (56) |
| CBS-2874 | S.Tm SL1344 | AmpR | WISH-5 |  | (56) |
| CBS-2875 | S.Tm SL1344 | AmpR | WISH-6 |  | (56) |
| CBS-2876 | S.Tm SL1344 | AmpR | WISH-8 |  | (56) |
| CBS-2877 | S.Tm SL1344 | AmpR | WISH-10 |  | (56) |
| CBS-1627 | S.Tm SL1344 | AmpR, CmR | disrupted sifA, sseJ, sseL (BE) | SPC914,p<br>SG089 | This study |
| CBS-1378 | S.Tm SL1344 | CmR | disrupted sifA, sseJ, sseL (BE) | SPC914 | This study |
| CBS-3156 | S.Tm SL1344 | AmpR | WISH-1 disrupted sifA, sseJ, sseL (BE) | cleared | This study |
| CBS-3155 | S.Tm SL1344 | AmpR | WISH-3 disrupted sifA, sseJ, sseL (BE) | cleared | This study |
| CBS-3152 | S.Tm SL1344 | AmpR | WISH-4 disrupted sifA, sseJ, sseL (BE) | cleared | This study |
| CBS-3154 | S.Tm SL1344 | AmpR | WISH-5 disrupted sifA, sseJ, sseL (BE) | cleared | This study |
| CBS-3151 | S.Tm SL1344 | AmpR | WISH-6 disrupted sifA, sseJ, sseL (BE) | cleared | This study |
| CBS-3153 | S.Tm SL1344 | AmpR | WISH-8 disrupted sifA, sseJ, sseL (BE) | cleared | This study |
| CBS-3084 | S.Tm SL1344 | AmpR | WISH-10 disrupted sifA, sseJ, sseL (BE) | cleared | This study |
| CBS-2499 | S.Tm SL1344 | AmpR | SL1344 disrupted sifB, srfJ, srgE, pipB,<br>cigR, steB (BE) | cleared | This study |
| CBS-1415 | S.Tm SL1344 | CmR | SL1344 disrupted sseF, sseG, sopD2,<br>steC, sifA, sseJ, gtgE, sspH2, pipB2 (BE) | SPC914 | This study |

**Table S2.**
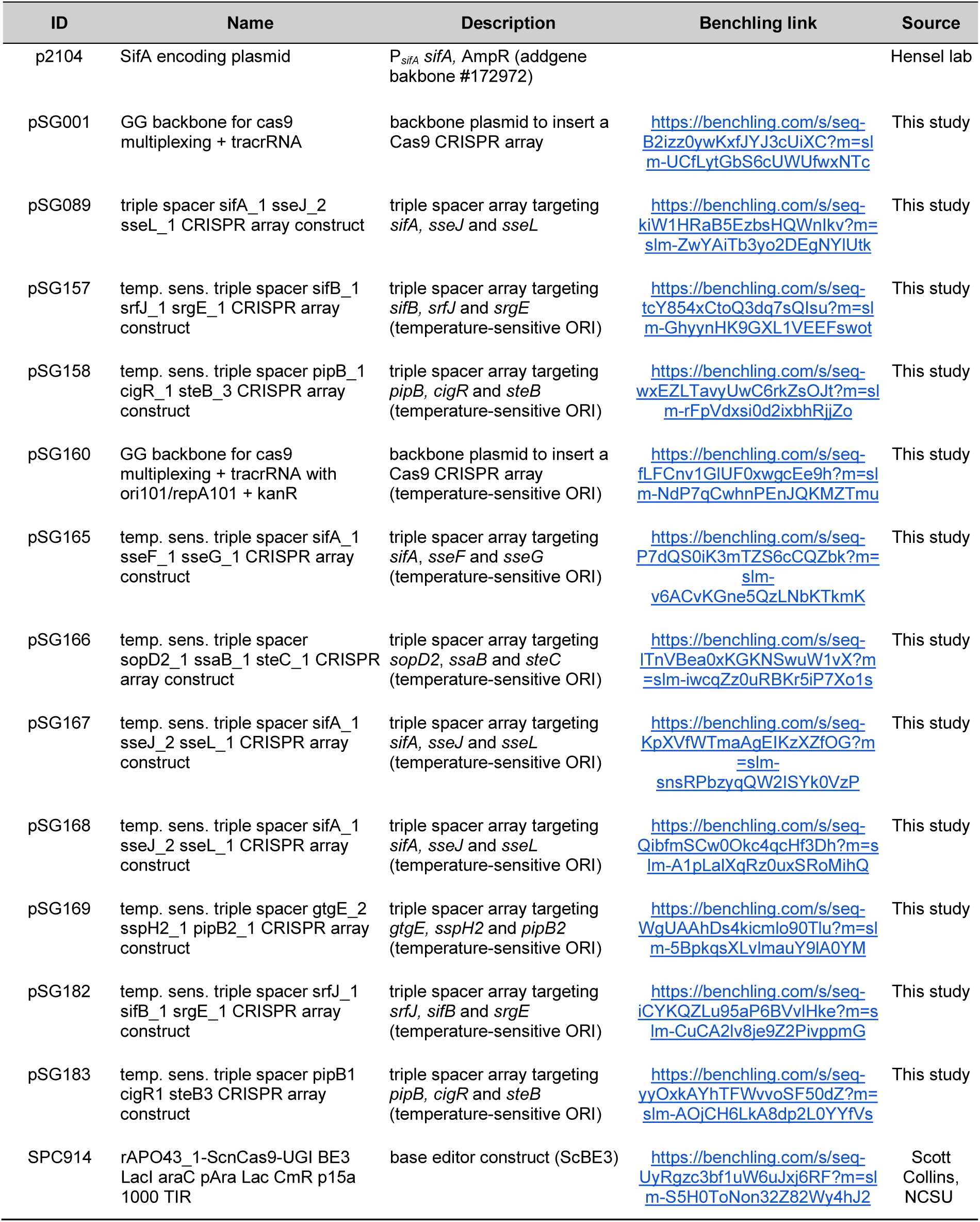
List of plasmids.

**Table S3.**
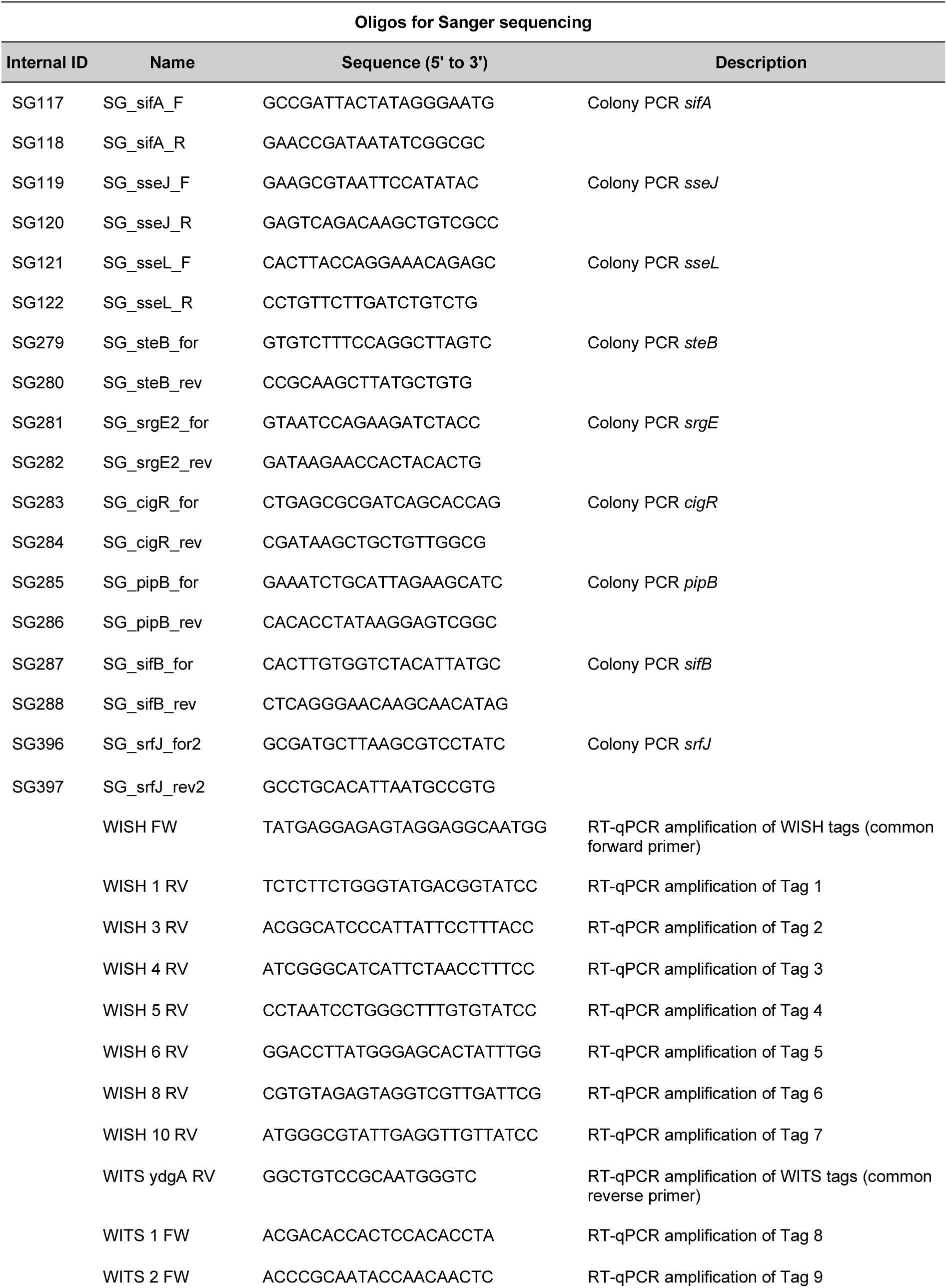
List of oligos.

**Table S4.**
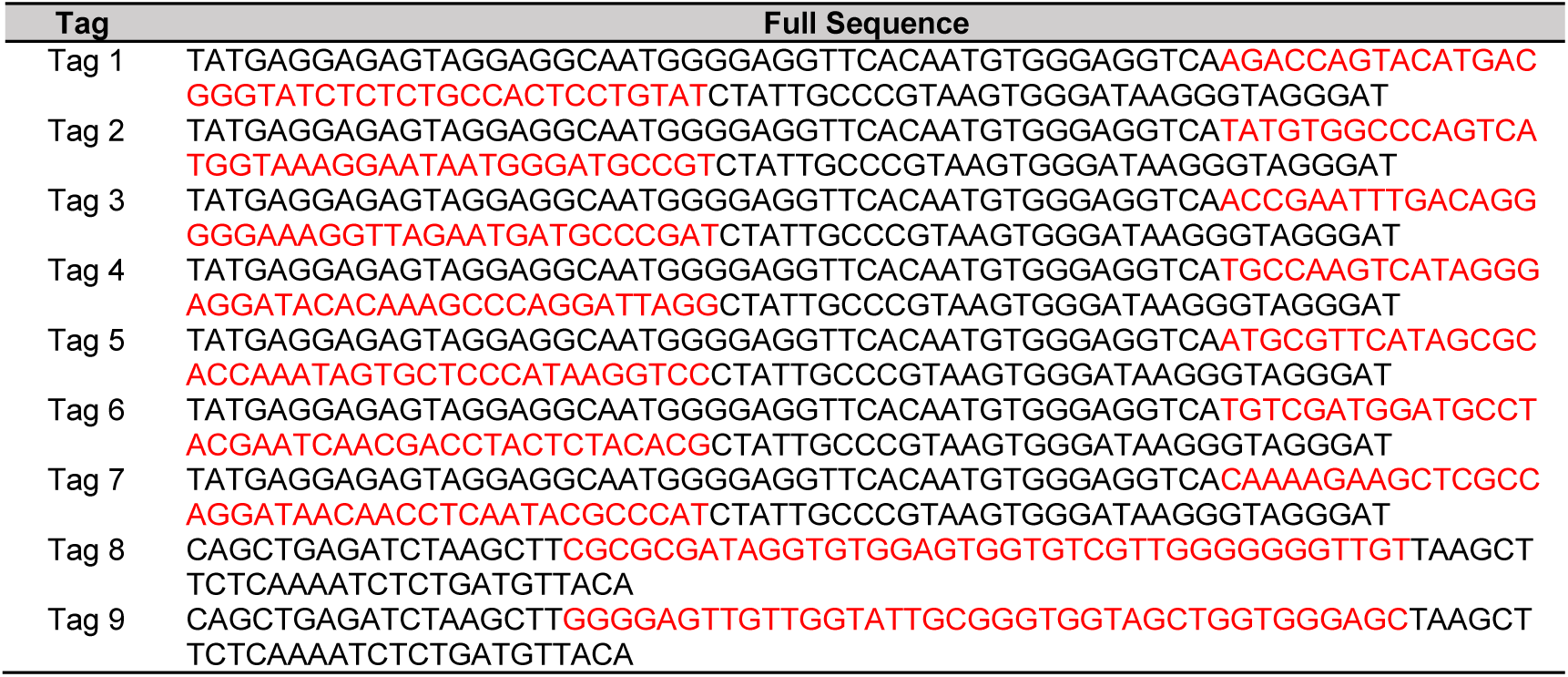
List of WISH-tag and WITS-tag sequences integrated in the genome of Salmonella enterica subsp. enterica serovar Typhimurium str. SL1344.

**Table S5.** List of SNVs for recombineered *sifA sseJ sseL*.

| Change | Minimum | Polymorphism Type | Coverage | Variant Frequency | Variant Raw Frequency | Average Quality |
| --- | --- | --- | --- | --- | --- | --- |
| *sifA, sseJ, sseL* clone 1 |  |  |  |  |  |  |
| C -> T | 4102464 | SNP (transition) | 61 | 44.26% | 185 | 24.93 |
| *sifA, sseJ, sseL* clone 2 |  |  |  |  |  |  |
| T -> C | 2381185 | SNP (transition) | 218 | 100% | 215 | 39.64 |

**Table S6.** List of SNVs for triple- and 6-fold *S*.Tm mutants.

| Change | Minimum | Polymorphism Type | Coverage | Variant Frequency | Variant Raw Frequency | Average Quality |
| --- | --- | --- | --- | --- | --- | --- |
| **CBS1378 triple-mutant *sifA*, *sseJ*, *sseL*** |  |  |  |  |  |  |
| C -> T | 1678387 | SNP (transition) | 186 | 99.5% | 185 | 32 |
| C -> T | 1678389 | SNP (transition) | 184 | 98.9% | 182 | 32 |
| C -> T | 2392562 | SNP (transition) | 177 | 100.0% | 177 | 32 |
| G -> A | 1266744 | SNP (transition) | 165 | 99.4% | 164 | 34 |
| C -> T | 1344780 | SNP (transition) | 194 | 100.0% | 194 | 34 |
| C -> T | 1534771 | SNP (transition) | 211 | 100.0% | 211 | 34 |
| C -> T | 1643207 | SNP (transition) | 220 | 99.5% | 219 | 34 |
| G -> A | 3399972 | SNP (transition) | 207 | 99.5% | 206 | 34 |
| G -> A | 964906 | SNP (transition) | 197 | 99.5% | 196 | 35 |
| C -> T | 1277548 | SNP (transition) | 201 | 100.0% | 201 | 35 |
| C -> T | 1428605 | SNP (transition) | 220 | 98.6% | 217 | 35 |
| G -> A | 1756092 | SNP (transition) | 155 | 100.0% | 155 | 35 |
| G -> A | 4595279 | SNP (transition) | 190 | 99.5% | 189 | 35 |
| T -> A | 635606 | SNP (transversion) | 226 | 100.0% | 226 | 36 |
| G -> A | 1687518 | SNP (transition) | 191 | 100.0% | 191 | 36 |
| G -> A | 1692794 | SNP (transition) | 193 | 100.0% | 193 | 36 |
| T -> C | 2411272 | SNP (transition) | 205 | 100.0% | 205 | 36 |
| **CBS-2499 6-fold mutant *sifB*, *srfJ*, *srgE*, *pipB*, *cigR*, *steB*** |  |  |  |  |  |  |
| C -> T | 829230 | SNP (transition) | 245 | 98.8% | 242 | 30 |
| C -> T | 982559 | SNP (transition) | 281 | 99.6% | 280 | 32 |
| G -> A | 1776327 | SNP (transition) | 212 | 100.0% | 212 | 32 |
| C -> T | 4689629 | SNP (transition) | 244 | 99.6% | 243 | 32 |
| C -> T | 111863 | SNP (transition) | 269 | 100.0% | 269 | 33 |
| C -> T | 881436 | SNP (transition) | 244 | 100.0% | 244 | 33 |
| C -> T | 1512358 | SNP (transition) | 220 | 100.0% | 220 | 33 |
| G -> A | 1890780 | SNP (transition) | 231 | 99.6% | 230 | 33 |
| G -> A | 2262539 | SNP (transition) | 228 | 100.0% | 228 | 33 |
| G -> A | 3201028 | SNP (transition) | 247 | 99.6% | 246 | 33 |
| C -> T | 4477145 | SNP (transition) | 252 | 100.0% | 252 | 33 |
| C -> T | 1002254 | SNP (transition) | 277 | 99.3% | 275 | 34 |
| G -> A | 2093565 | SNP (transition) | 209 | 99.5% | 208 | 34 |
| G -> A | 2617708 | SNP (transition) | 262 | 100.0% | 262 | 34 |
| G -> A | 2671789 | SNP (transition) | 217 | 100.0% | 217 | 34 |
| G -> A | 3140875 | SNP (transition) | 251 | 99.6% | 250 | 34 |
| C -> T | 4395554 | SNP (transition) | 270 | 100.0% | 270 | 34 |
| C -> T | 4567578 | SNP (transition) | 279 | 99.6% | 278 | 34 |
| C -> T | 4715079 | SNP (transition) | 253 | 99.6% | 252 | 34 |
| C -> T | 433851 | SNP (transition) | 214 | 100.0% | 214 | 35 |
| C -> T | 783952 | SNP (transition) | 221 | 100.0% | 221 | 35 |
| C -> T | 1331325 | SNP (transition) | 273 | 100.0% | 273 | 35 |
| C -> T | 1515888 | SNP (transition) | 252 | 100.0% | 252 | 35 |
| G -> A | 1588565 | SNP (transition) | 225 | 99.6% | 224 | 35 |
| C -> T | 1649093 | SNP (transition) | 221 | 98.6% | 218 | 35 |
| C -> T | 1649097 | SNP (transition) | 224 | 99.6% | 223 | 35 |
| G -> A | 1676476 | SNP (transition) | 239 | 99.6% | 238 | 35 |
| G -> A | 1756092 | SNP (transition) | 236 | 99.6% | 235 | 35 |
| G -> A | 3454457 | SNP (transition) | 231 | 100.0% | 231 | 35 |
| G -> A | 3494576 | SNP (transition) | 231 | 99.6% | 230 | 35 |
| C -> T | 3982539 | SNP (transition) | 236 | 100.0% | 236 | 35 |
| G -> A | 4061474 | SNP (transition) | 291 | 100.0% | 291 | 35 |
| C -> T | 4356150 | SNP (transition) | 264 | 99.6% | 263 | 35 |
| C -> T | 4624425 | SNP (transition) | 264 | 100.0% | 264 | 35 |
| G -> A | 4648015 | SNP (transition) | 229 | 99.1% | 227 | 35 |
| C -> T | 4683220 | SNP (transition) | 238 | 99.6% | 237 | 35 |
| C -> T | 4689632 | SNP (transition) | 239 | 100.0% | 239 | 35 |
| C -> T | 4751082 | SNP (transition) | 272 | 100.0% | 272 | 35 |
| C -> T | 4876234 | SNP (transition) | 212 | 99.1% | 210 | 35 |
| T -> A | 635606 | SNP (transversion) | 221 | 98.6% | 218 | 36 |
| C -> T | 1133535 | SNP (transition) | 212 | 100.0% | 212 | 36 |
| C -> T | 1308769 | SNP (transition) | 192 | 100.0% | 192 | 36 |
| C -> T | 1382265 | SNP (transition) | 207 | 100.0% | 207 | 36 |
| G -> A | 3720748 | SNP (transition) | 284 | 100.0% | 284 | 36 |
| T -> C | 2411272 | SNP (transition) | 230 | 99.6% | 229 | 37 |

**Table S7.** List of SNVs for WISH-tagged *sifA, sseJ, sseL* mutants.

| Change | Minimum | Polymorphism Type | Coverage | Variant Frequency | Variant Raw Frequency | Average Quality |
| --- | --- | --- | --- | --- | --- | --- |
| **SG493 *sifA*, *sseJ*, *sseL* WISH-1 JN1** |  |  |  |  |  |  |
| C -> T | 25617 | SNP (transition) | 271 | 100.0% | 271 | 35 |
| C -> T | 58280 | SNP (transition) | 260 | 100.0% | 260 | 36 |
| C -> T | 68805 | SNP (transition) | 260 | 99.6% | 259 | 36 |
| C -> T | 90656 | SNP (transition) | 186 | 99.5% | 185 | 35 |
| C -> T | 119748 | SNP (transition) | 223 | 99.6% | 222 | 35 |
| C -> T | 127625 | SNP (transition) | 335 | 99.7% | 334 | 34 |
| C -> T | 132512 | SNP (transition) | 299 | 99.3% | 297 | 35 |
| C -> T | 136579 | SNP (transition) | 239 | 99.6% | 238 | 36 |
| C -> T | 141931 | SNP (transition) | 188 | 100.0% | 188 | 35 |
| C -> T | 148772 | SNP (transition) | 235 | 100.0% | 235 | 35 |
| C -> T | 170191 | SNP (transition) | 243 | 100.0% | 243 | 36 |
| C -> T | 171394 | SNP (transition) | 207 | 100.0% | 207 | 36 |
| C -> T | 171460 | SNP (transition) | 217 | 100.0% | 217 | 36 |
| C -> T | 280345 | SNP (transition) | 249 | 100.0% | 249 | 35 |
| C -> T | 297601 | SNP (transition) | 251 | 99.2% | 249 | 33 |
| C -> T | 338082 | SNP (transition) | 232 | 100.0% | 232 | 35 |
| T -> A | 635606 | SNP (transversion) | 264 | 100.0% | 264 | 36 |
| C -> T | 727319 | SNP (transition) | 247 | 100.0% | 247 | 33 |
| C -> T | 915656 | SNP (transition) | 281 | 100.0% | 281 | 35 |
| G -> A | 972236 | SNP (transition) | 219 | 100.0% | 219 | 34 |
| C -> T | 1123580 | SNP (transition) | 226 | 99.6% | 225 | 35 |
| C -> T | 1134482 | SNP (transition) | 243 | 99.6% | 242 | 36 |
| G -> A | 1266744 | SNP (transition) | 229 | 100.0% | 229 | 35 |
| C -> T | 1633603 | SNP (transition) | 278 | 99.6% | 277 | 35 |
| C -> T | 1637646 | SNP (transition) | 244 | 100.0% | 244 | 36 |
| G -> A | 1639151 | SNP (transition) | 235 | 100.0% | 235 | 35 |
| G -> A | 1672670 | SNP (transition) | 213 | 100.0% | 213 | 35 |
| C -> T | 1675780 | SNP (transition) | 227 | 100.0% | 227 | 35 |
| C -> T | 1678387 | SNP (transition) | 255 | 100.0% | 255 | 34 |
| C -> T | 1678389 | SNP (transition) | 249 | 99.6% | 248 | 35 |
| G -> A | 1711196 | SNP (transition) | 226 | 100.0% | 226 | 36 |
| G -> A | 1756092 | SNP (transition) | 215 | 100.0% | 215 | 36 |
| G -> A | 1813320 | SNP (transition) | 230 | 100.0% | 230 | 36 |
| G -> A | 1886775 | SNP (transition) | 203 | 100.0% | 203 | 34 |
| G -> A | 1947607 | SNP (transition) | 240 | 100.0% | 240 | 35 |
| G -> A | 2026302 | SNP (transition) | 224 | 100.0% | 224 | 36 |
| G -> A | 2066341 | SNP (transition) | 208 | 100.0% | 208 | 35 |
| G -> A | 2117007 | SNP (transition) | 249 | 100.0% | 249 | 36 |
| G -> A | 2139160 | SNP (transition) | 243 | 100.0% | 243 | 35 |
| G -> A | 2167707 | SNP (transition) | 268 | 100.0% | 268 | 36 |
| G -> A | 2170378 | SNP (transition) | 202 | 100.0% | 202 | 36 |
| G -> A | 2171431 | SNP (transition) | 242 | 100.0% | 242 | 35 |
| G -> A | 2183015 | SNP (transition) | 229 | 99.6% | 228 | 35 |
| G -> A | 2185186 | SNP (transition) | 232 | 99.6% | 231 | 35 |
| G -> A | 2186271 | SNP (transition) | 266 | 100.0% | 266 | 34 |
| G -> A | 2193451 | SNP (transition) | 226 | 100.0% | 226 | 35 |
| G -> A | 2195473 | SNP (transition) | 234 | 98.7% | 231 | 32 |
| G -> A | 2197463 | SNP (transition) | 258 | 99.6% | 257 | 35 |
| G -> A | 2209798 | SNP (transition) | 263 | 100.0% | 263 | 36 |
| G -> A | 2220590 | SNP (transition) | 193 | 100.0% | 193 | 35 |
| G -> A | 2220924 | SNP (transition) | 215 | 100.0% | 215 | 34 |
| G -> A | 2225243 | SNP (transition) | 267 | 100.0% | 267 | 37 |
| G -> A | 2231765 | SNP (transition) | 242 | 99.6% | 241 | 36 |
| G -> A | 2235543 | SNP (transition) | 282 | 99.6% | 281 | 35 |
| G -> A | 2236886 | SNP (transition) | 236 | 100.0% | 236 | 35 |
| G -> A | 2236970 | SNP (transition) | 244 | 100.0% | 244 | 34 |
| G -> A | 2239831 | SNP (transition) | 257 | 100.0% | 257 | 35 |
| G -> A | 2239926 | SNP (transition) | 265 | 100.0% | 265 | 36 |
| G -> A | 2248607 | SNP (transition) | 259 | 100.0% | 259 | 36 |
| G -> A | 2253524 | SNP (transition) | 235 | 99.6% | 234 | 36 |
| G -> A | 2258249 | SNP (transition) | 211 | 100.0% | 211 | 36 |
| G -> A | 2268011 | SNP (transition) | 238 | 99.6% | 237 | 35 |
| G -> A | 2269616 | SNP (transition) | 231 | 99.6% | 230 | 37 |
| G -> A | 2271630 | SNP (transition) | 276 | 100.0% | 276 | 35 |
| G -> A | 2273663 | SNP (transition) | 227 | 99.6% | 226 | 36 |
| G -> A | 2281283 | SNP (transition) | 238 | 100.0% | 238 | 36 |
| G -> A | 2294675 | SNP (transition) | 257 | 100.0% | 257 | 34 |
| G -> A | 2324400 | SNP (transition) | 259 | 99.6% | 258 | 34 |
| G -> A | 2326156 | SNP (transition) | 209 | 100.0% | 209 | 35 |
| G -> A | 2359657 | SNP (transition) | 194 | 100.0% | 194 | 36 |
| G -> A | 2378522 | SNP (transition) | 256 | 100.0% | 256 | 37 |
| C -> T | 2392562 | SNP (transition) | 203 | 99.5% | 202 | 34 |
| T -> C | 2411272 | SNP (transition) | 236 | 100.0% | 236 | 36 |
| G -> A | 2486514 | SNP (transition) | 223 | 99.6% | 222 | 36 |
| G -> A | 2522113 | SNP (transition) | 221 | 100.0% | 221 | 34 |
| C -> T | 2529542 | SNP (transition) | 231 | 100.0% | 231 | 35 |
| G -> A | 2637243 | SNP (transition) | 227 | 100.0% | 227 | 35 |
| G -> A | 2776670 | SNP (transition) | 248 | 100.0% | 248 | 34 |
| G -> A | 2781250 | SNP (transition) | 250 | 100.0% | 250 | 35 |
| G -> A | 2805136 | SNP (transition) | 246 | 99.2% | 244 | 35 |
| C -> T | 3009946 | SNP (transition) | 217 | 99.5% | 216 | 34 |
| G -> A | 3237765 | SNP (transition) | 242 | 100.0% | 242 | 36 |
| C -> T | 3241806 | SNP (transition) | 215 | 100.0% | 215 | 34 |
| G -> A | 3261826 | SNP (transition) | 223 | 100.0% | 223 | 33 |
| C -> T | 3309182 | SNP (transition) | 252 | 100.0% | 252 | 36 |
| G -> A | 3412901 | SNP (transition) | 286 | 99.7% | 285 | 32 |
| G -> A | 3435522 | SNP (transition) | 248 | 98.4% | 244 | 33 |
| G -> A | 3535514 | SNP (transition) | 279 | 100.0% | 279 | 35 |
| C -> T | 3670583 | SNP (transition) | 306 | 99.7% | 305 | 35 |
| G -> A | 3815931 | SNP (transition) | 229 | 100.0% | 229 | 34 |
| G -> A | 3975555 | SNP (transition) | 252 | 99.6% | 251 | 36 |
| C -> T | 4171568 | SNP (transition) | 259 | 100.0% | 259 | 36 |
| C -> T | 4186275 | SNP (transition) | 256 | 100.0% | 256 | 35 |
| C -> T | 4199400 | SNP (transition) | 255 | 100.0% | 255 | 36 |
| C -> T | 4245941 | SNP (transition) | 224 | 100.0% | 224 | 36 |
| C -> T | 4250357 | SNP (transition) | 281 | 99.3% | 279 | 35 |
| C -> T | 4321145 | SNP (transition) | 309 | 100.0% | 309 | 35 |
| C -> T | 4334183 | SNP (transition) | 252 | 100.0% | 252 | 35 |
| G -> A | 4368682 | SNP (transition) | 246 | 99.2% | 244 | 33 |
| C -> T | 4389707 | SNP (transition) | 282 | 100.0% | 282 | 36 |
| C -> T | 4398050 | SNP (transition) | 206 | 99.5% | 205 | 35 |
| C -> T | 4405896 | SNP (transition) | 228 | 100.0% | 228 | 36 |
| C -> T | 4415013 | SNP (transition) | 225 | 100.0% | 225 | 36 |
| C -> T | 4438810 | SNP (transition) | 247 | 99.6% | 246 | 36 |
| C -> T | 4442827 | SNP (transition) | 240 | 98.3% | 236 | 33 |
| C -> T | 4445350 | SNP (transition) | 238 | 100.0% | 238 | 36 |
| C -> T | 4451185 | SNP (transition) | 344 | 99.1% | 341 | 35 |
| C -> T | 4453247 | SNP (transition) | 292 | 100.0% | 292 | 36 |
| C -> T | 4460644 | SNP (transition) | 237 | 100.0% | 237 | 36 |
| C -> T | 4473624 | SNP (transition) | 260 | 99.6% | 259 | 36 |
| C -> T | 4494958 | SNP (transition) | 315 | 100.0% | 315 | 35 |
| C -> T | 4499408 | SNP (transition) | 246 | 99.6% | 245 | 36 |
| C -> T | 4500667 | SNP (transition) | 223 | 100.0% | 223 | 36 |
| C -> T | 4500870 | SNP (transition) | 275 | 100.0% | 275 | 35 |
| C -> T | 4503857 | SNP (transition) | 281 | 100.0% | 281 | 35 |
| C -> T | 4507151 | SNP (transition) | 254 | 99.6% | 253 | 36 |
| C -> T | 4591060 | SNP (transition) | 249 | 99.6% | 248 | 36 |
| C -> T | 4597994 | SNP (transition) | 256 | 100.0% | 256 | 35 |
| C -> T | 4681759 | SNP (transition) | 272 | 100.0% | 272 | 34 |
| C -> T | 4693779 | SNP (transition) | 214 | 99.5% | 213 | 34 |
| C -> T | 4745632 | SNP (transition) | 282 | 99.6% | 281 | 34 |
| T -> C | 4790592 | SNP (transition) | 249 | 100.0% | 249 | 37 |
| C -> T | 4808935 | SNP (transition) | 253 | 100.0% | 253 | 36 |
| **SG492 *sifA*, *sseJ*, *sseL* WISH-3 JN2** |  |  |  |  |  |  |
| G -> A | 4831565 | SNP (transition) | 320 | 99.7% | 319 | 34 |
| C -> T | 251356 | SNP (transition) | 264 | 99.6% | 263 | 35 |
| C -> T | 407652 | SNP (transition) | 327 | 100.0% | 327 | 35 |
| T -> A | 635606 | SNP (transversion) | 256 | 100.0% | 256 | 36 |
| G -> A | 697841 | SNP (transition) | 248 | 99.2% | 246 | 32 |
| A -> G | 924260 | SNP (transition) | 239 | 89.5% | 214 | 31 |
| C -> T | 1054715 | SNP (transition) | 252 | 99.6% | 251 | 35 |
| G -> A | 1266744 | SNP (transition) | 268 | 100.0% | 268 | 36 |
| C -> T | 1364531 | SNP (transition) | 276 | 100.0% | 276 | 36 |
| C -> T | 1429776 | SNP (transition) | 238 | 100.0% | 238 | 35 |
| C -> T | 1483116 | SNP (transition) | 215 | 99.5% | 214 | 34 |
| C -> T | 1514475 | SNP (transition) | 260 | 100.0% | 260 | 36 |
| C -> T | 1678384 | SNP (transition) | 216 | 100.0% | 216 | 36 |
| C -> T | 1678387 | SNP (transition) | 205 | 100.0% | 205 | 34 |
| C -> T | 1678389 | SNP (transition) | 200 | 100.0% | 200 | 35 |
| C -> T | 1680450 | SNP (transition) | 246 | 99.6% | 245 | 36 |
| G -> A | 1732335 | SNP (transition) | 224 | 100.0% | 224 | 33 |
| G -> A | 1756092 | SNP (transition) | 257 | 99.2% | 255 | 36 |
| G -> A | 1975083 | SNP (transition) | 231 | 100.0% | 231 | 33 |
| G -> A | 1995372 | SNP (transition) | 303 | 99.7% | 302 | 35 |
| G -> A | 2076967 | SNP (transition) | 220 | 100.0% | 220 | 35 |
| G -> A | 2091978 | SNP (transition) | 251 | 98.8% | 248 | 36 |
| G -> A | 2162998 | SNP (transition) | 257 | 100.0% | 257 | 36 |
| G -> A | 2195469 | SNP (transition) | 256 | 99.6% | 255 | 33 |
| C -> T | 2205937 | SNP (transition) | 258 | 99.2% | 256 | 33 |
| G -> A | 2233791 | SNP (transition) | 250 | 99.6% | 249 | 34 |
| G -> A | 2253060 | SNP (transition) | 249 | 100.0% | 249 | 36 |
| G -> A | 2312812 | SNP (transition) | 251 | 100.0% | 251 | 35 |
| C -> T | 2392562 | SNP (transition) | 239 | 100.0% | 239 | 35 |
| T -> C | 2411272 | SNP (transition) | 236 | 100.0% | 236 | 36 |
| G -> A | 2532291 | SNP (transition) | 275 | 100.0% | 275 | 35 |
| G -> A | 2811509 | SNP (transition) | 278 | 100.0% | 278 | 35 |
| G -> A | 2824398 | SNP (transition) | 307 | 99.3% | 305 | 28 |
| G -> A | 3078336 | SNP (transition) | 209 | 100.0% | 209 | 35 |
| G -> A | 3489969 | SNP (transition) | 309 | 99.7% | 308 | 34 |
| G -> A | 3643086 | SNP (transition) | 313 | 100.0% | 313 | 35 |
| G -> A | 3647753 | SNP (transition) | 325 | 100.0% | 325 | 35 |
| G -> A | 3772713 | SNP (transition) | 293 | 100.0% | 293 | 35 |
| G -> A | 3875355 | SNP (transition) | 254 | 100.0% | 254 | 37 |
| G -> A | 3927934 | SNP (transition) | 343 | 100.0% | 343 | 33 |
| G -> A | 3949175 | SNP (transition) | 335 | 99.4% | 333 | 36 |
| G -> A | 4101952 | SNP (transition) | 241 | 100.0% | 241 | 36 |
| C -> T | 4105437 | SNP (transition) | 236 | 99.6% | 235 | 36 |
| C -> T | 4294190 | SNP (transition) | 277 | 100.0% | 277 | 36 |
| C -> T | 4319589 | SNP (transition) | 262 | 99.6% | 261 | 36 |
| G -> A | 4368682 | SNP (transition) | 324 | 100.0% | 324 | 32 |
| C -> T | 4369446 | SNP (transition) | 272 | 100.0% | 272 | 36 |
| C -> T | 4524783 | SNP (transition) | 292 | 100.0% | 292 | 36 |
| C -> T | 4591655 | SNP (transition) | 296 | 99.0% | 293 | 35 |
| C -> T | 4596962 | SNP (transition) | 256 | 100.0% | 256 | 36 |
| C -> T | 4609924 | SNP (transition) | 214 | 100.0% | 214 | 35 |
| **SG489 *sifA*, *sseJ*, *sseL* WISH-4 JN5** |  |  |  |  |  |  |
| C -> T | 83067 | SNP (transition) | 333 | 99.7% | 332 | 34 |
| C -> T | 229597 | SNP (transition) | 286 | 100.0% | 286 | 34 |
| C -> T | 229783 | SNP (transition) | 210 | 100.0% | 210 | 34 |
| C -> T | 239793 | SNP (transition) | 294 | 100.0% | 294 | 35 |
| C -> T | 284638 | SNP (transition) | 321 | 100.0% | 321 | 34 |
| C -> T | 306315 | SNP (transition) | 303 | 100.0% | 303 | 34 |
| C -> T | 353695 | SNP (transition) | 312 | 99.7% | 311 | 36 |
| C -> T | 402695 | SNP (transition) | 302 | 99.3% | 300 | 29 |
| C -> T | 435779 | SNP (transition) | 315 | 99.7% | 314 | 35 |
| C -> T | 457533 | SNP (transition) | 324 | 100.0% | 324 | 35 |
| C -> T | 467583 | SNP (transition) | 287 | 100.0% | 287 | 36 |
| C -> T | 472008 | SNP (transition) | 329 | 99.4% | 327 | 35 |
| C -> T | 474064 | SNP (transition) | 304 | 100.0% | 304 | 36 |
| C -> T | 497943 | SNP (transition) | 280 | 100.0% | 280 | 34 |
| C -> T | 507982 | SNP (transition) | 282 | 99.6% | 281 | 35 |
| C -> T | 520672 | SNP (transition) | 345 | 99.7% | 344 | 34 |
| T -> A | 635606 | SNP (transversion) | 374 | 100.0% | 374 | 36 |
| C -> T | 1226916 | SNP (transition) | 272 | 100.0% | 272 | 33 |
| G -> A | 1266744 | SNP (transition) | 253 | 100.0% | 253 | 35 |
| C -> T | 1678387 | SNP (transition) | 281 | 99.3% | 279 | 33 |
| C -> T | 1678389 | SNP (transition) | 275 | 100.0% | 275 | 34 |
| G -> A | 1756092 | SNP (transition) | 243 | 100.0% | 243 | 35 |
| G -> A | 2369679 | SNP (transition) | 262 | 100.0% | 262 | 36 |
| C -> T | 2392562 | SNP (transition) | 259 | 99.6% | 258 | 34 |
| T -> C | 2411272 | SNP (transition) | 341 | 100.0% | 341 | 36 |
| G -> A | 2437817 | SNP (transition) | 264 | 100.0% | 264 | 34 |
| G -> A | 3084418 | SNP (transition) | 278 | 99.3% | 276 | 34 |
| G -> A | 3508561 | SNP (transition) | 336 | 99.7% | 335 | 35 |
| C -> T | 4207714 | SNP (transition) | 335 | 100.0% | 335 | 36 |
| C -> T | 4391871 | SNP (transition) | 318 | 99.7% | 317 | 35 |
| C -> T | 4591655 | SNP (transition) | 304 | 99.0% | 301 | 33 |
| C -> T | 4683109 | SNP (transition) | 309 | 99.7% | 308 | 36 |
| C -> T | 4818137 | SNP (transition) | 304 | 99.7% | 303 | 35 |
| C -> T | 4818414 | SNP (transition) | 346 | 100.0% | 346 | 36 |
| C -> T | 4820794 | SNP (transition) | 297 | 100.0% | 297 | 37 |
| **SG491 *sifA*, *sseJ*, *sseL* WISH-5 JN6** |  |  |  |  |  |  |
| C -> T | 90155 | SNP (transition) | 340 | 100.0% | 340 | 35 |
| C -> T | 118499 | SNP (transition) | 306 | 100.0% | 306 | 34 |
| C -> T | 186959 | SNP (transition) | 277 | 98.9% | 274 | 35 |
| C -> T | 230023 | SNP (transition) | 77 | 100.0% | 77 | 34 |
| C -> T | 280497 | SNP (transition) | 238 | 100.0% | 238 | 33 |
| C -> T | 329717 | SNP (transition) | 150 | 99.3% | 149 | 34 |
| C -> T | 410917 | SNP (transition) | 236 | 100.0% | 236 | 32 |
| C -> T | 439825 | SNP (transition) | 294 | 100.0% | 294 | 34 |
| C -> T | 485834 | SNP (transition) | 257 | 100.0% | 257 | 32 |
| C -> T | 489055 | SNP (transition) | 274 | 100.0% | 274 | 36 |
| C -> T | 492825 | SNP (transition) | 166 | 99.4% | 165 | 33 |
| C -> T | 514102 | SNP (transition) | 281 | 100.0% | 281 | 34 |
| C -> T | 572270 | SNP (transition) | 274 | 99.3% | 272 | 32 |
| C -> T | 576573 | SNP (transition) | 291 | 99.7% | 290 | 35 |
| C -> T | 583222 | SNP (transition) | 230 | 100.0% | 230 | 35 |
| C -> T | 624779 | SNP (transition) | 256 | 100.0% | 256 | 36 |
| T -> A | 635606 | SNP (transversion) | 297 | 99.7% | 296 | 36 |
| C -> T | 647020 | SNP (transition) | 251 | 99.2% | 249 | 34 |
| C -> T | 656855 | SNP (transition) | 311 | 100.0% | 311 | 34 |
| C -> T | 657370 | SNP (transition) | 292 | 99.0% | 289 | 34 |
| C -> T | 668032 | SNP (transition) | 187 | 100.0% | 187 | 35 |
| C -> T | 899342 | SNP (transition) | 221 | 100.0% | 221 | 32 |
| C -> T | 944314 | SNP (transition) | 196 | 99.5% | 195 | 32 |
| C -> T | 1146131 | SNP (transition) | 303 | 100.0% | 303 | 33 |
| G -> A | 1266744 | SNP (transition) | 163 | 100.0% | 163 | 36 |
| C -> T | 1465320 | SNP (transition) | 244 | 100.0% | 244 | 34 |
| C -> T | 1535575 | SNP (transition) | 253 | 100.0% | 253 | 34 |
| C -> T | 1552020 | SNP (transition) | 293 | 100.0% | 293 | 35 |
| G -> A | 1607477 | SNP (transition) | 275 | 99.6% | 274 | 35 |
| C -> T | 1637376 | SNP (transition) | 221 | 100.0% | 221 | 35 |
| C -> T | 1651424 | SNP (transition) | 255 | 100.0% | 255 | 34 |
| C -> T | 1666047 | SNP (transition) | 192 | 100.0% | 192 | 35 |
| C -> T | 1678387 | SNP (transition) | 172 | 99.4% | 171 | 34 |
| C -> T | 1678389 | SNP (transition) | 175 | 100.0% | 175 | 35 |
| C -> T | 1687015 | SNP (transition) | 206 | 99.5% | 205 | 35 |
| C -> T | 1691549 | SNP (transition) | 263 | 100.0% | 263 | 36 |
| G -> A | 1756092 | SNP (transition) | 239 | 99.6% | 238 | 36 |
| G -> A | 1868087 | SNP (transition) | 299 | 100.0% | 299 | 34 |
| G -> A | 1933302 | SNP (transition) | 205 | 100.0% | 205 | 35 |
| G -> A | 1959609 | SNP (transition) | 305 | 100.0% | 305 | 34 |
| G -> A | 1986757 | SNP (transition) | 137 | 98.5% | 135 | 34 |
| G -> A | 1987576 | SNP (transition) | 269 | 100.0% | 269 | 34 |
| G -> A | 2007912 | SNP (transition) | 285 | 99.6% | 284 | 33 |
| G -> A | 2048002 | SNP (transition) | 273 | 100.0% | 273 | 36 |
| G -> A | 2281045 | SNP (transition) | 282 | 100.0% | 282 | 35 |
| G -> A | 2282676 | SNP (transition) | 195 | 98.5% | 192 | 34 |
| C -> T | 2392562 | SNP (transition) | 161 | 100.0% | 161 | 34 |
| T -> C | 2411272 | SNP (transition) | 304 | 100.0% | 304 | 36 |
| G -> A | 2415412 | SNP (transition) | 252 | 100.0% | 252 | 34 |
| G -> A | 2416462 | SNP (transition) | 250 | 99.6% | 249 | 33 |
| G -> A | 2419102 | SNP (transition) | 237 | 100.0% | 237 | 35 |
| G -> A | 2426821 | SNP (transition) | 301 | 99.7% | 300 | 35 |
| G -> A | 2452078 | SNP (transition) | 247 | 100.0% | 247 | 35 |
| G -> A | 2461180 | SNP (transition) | 263 | 99.6% | 262 | 35 |
| G -> A | 2509463 | SNP (transition) | 198 | 100.0% | 198 | 34 |
| G -> A | 2509919 | SNP (transition) | 264 | 100.0% | 264 | 35 |
| G -> A | 2671282 | SNP (transition) | 291 | 100.0% | 291 | 34 |
| G -> A | 2751233 | SNP (transition) | 331 | 100.0% | 331 | 34 |
| G -> A | 2772846 | SNP (transition) | 315 | 99.7% | 314 | 27 |
| G -> A | 2775664 | SNP (transition) | 220 | 73.2% | 161 | 24 |
| G -> A | 2787701 | SNP (transition) | 338 | 99.4% | 336 | 34 |
| G -> A | 2807603 | SNP (transition) | 218 | 100.0% | 218 | 33 |
| G -> A | 2832513 | SNP (transition) | 192 | 99.5% | 191 | 35 |
| G -> A | 2834228 | SNP (transition) | 254 | 99.6% | 253 | 36 |
| G -> A | 2834695 | SNP (transition) | 242 | 100.0% | 242 | 35 |
| G -> A | 2878360 | SNP (transition) | 207 | 100.0% | 207 | 34 |
| G -> A | 2883383 | SNP (transition) | 243 | 100.0% | 243 | 35 |
| G -> A | 2885395 | SNP (transition) | 244 | 99.2% | 242 | 34 |
| G -> A | 2894046 | SNP (transition) | 159 | 99.4% | 158 | 34 |
| G -> A | 2901525 | SNP (transition) | 321 | 100.0% | 321 | 34 |
| G -> A | 2923254 | SNP (transition) | 306 | 100.0% | 306 | 35 |
| G -> A | 2924217 | SNP (transition) | 134 | 100.0% | 134 | 36 |
| G -> A | 2934285 | SNP (transition) | 265 | 100.0% | 265 | 28 |
| G -> A | 2935951 | SNP (transition) | 229 | 99.6% | 228 | 35 |
| G -> A | 2949832 | SNP (transition) | 229 | 100.0% | 229 | 35 |
| G -> A | 2954155 | SNP (transition) | 258 | 100.0% | 258 | 35 |
| G -> A | 2992260 | SNP (transition) | 295 | 99.3% | 293 | 35 |
| G -> A | 3014050 | SNP (transition) | 308 | 100.0% | 308 | 35 |
| G -> A | 3042730 | SNP (transition) | 262 | 100.0% | 262 | 36 |
| G -> A | 3058897 | SNP (transition) | 217 | 99.5% | 216 | 33 |
| G -> A | 3065565 | SNP (transition) | 190 | 99.5% | 189 | 35 |
| G -> A | 3089795 | SNP (transition) | 299 | 99.7% | 298 | 35 |
| G -> A | 3093621 | SNP (transition) | 315 | 100.0% | 315 | 36 |
| G -> A | 3119025 | SNP (transition) | 242 | 100.0% | 242 | 35 |
| G -> A | 3133013 | SNP (transition) | 283 | 100.0% | 283 | 34 |
| G -> A | 3142297 | SNP (transition) | 274 | 100.0% | 274 | 34 |
| G -> A | 3163189 | SNP (transition) | 268 | 99.6% | 267 | 36 |
| G -> A | 3188091 | SNP (transition) | 266 | 99.6% | 265 | 34 |
| G -> A | 3213354 | SNP (transition) | 142 | 100.0% | 142 | 35 |
| G -> A | 3257062 | SNP (transition) | 199 | 99.5% | 198 | 32 |
| G -> A | 3348386 | SNP (transition) | 240 | 99.2% | 238 | 34 |
| G -> A | 3391070 | SNP (transition) | 295 | 99.7% | 294 | 34 |
| G -> A | 3395040 | SNP (transition) | 253 | 99.2% | 251 | 35 |
| G -> A | 3397900 | SNP (transition) | 263 | 100.0% | 263 | 33 |
| G -> A | 3400976 | SNP (transition) | 256 | 100.0% | 256 | 35 |
| G -> A | 3409512 | SNP (transition) | 250 | 99.6% | 249 | 34 |
| G -> A | 3515017 | SNP (transition) | 313 | 100.0% | 313 | 35 |
| G -> A | 3573372 | SNP (transition) | 263 | 100.0% | 263 | 35 |
| G -> A | 3587573 | SNP (transition) | 210 | 99.5% | 209 | 35 |
| G -> A | 3595148 | SNP (transition) | 230 | 100.0% | 230 | 35 |
| G -> A | 3604981 | SNP (transition) | 275 | 100.0% | 275 | 35 |
| C -> T | 3636949 | SNP (transition) | 275 | 99.6% | 274 | 36 |
| G -> A | 3735364 | SNP (transition) | 263 | 94.3% | 248 | 30 |
| G -> A | 3763231 | SNP (transition) | 250 | 100.0% | 250 | 34 |
| G -> A | 4058860 | SNP (transition) | 243 | 100.0% | 243 | 35 |
| G -> A | 4087266 | SNP (transition) | 205 | 100.0% | 205 | 36 |
| G -> A | 4261872 | SNP (transition) | 237 | 100.0% | 237 | 34 |
| G -> A | 4368682 | SNP (transition) | 298 | 99.3% | 296 | 32 |
| C -> T | 4714914 | SNP (transition) | 273 | 99.3% | 271 | 32 |
| C -> T | 4782544 | SNP (transition) | 201 | 100.0% | 201 | 37 |
| C -> T | 4826895 | SNP (transition) | 227 | 100.0% | 227 | 34 |
| **SG488 sifA, sseJ, sseL WISH-6 JN5** |  |  |  |  |  |  |
| G -> A | 57967 | SNP (transition) | 315 | 99.7% | 314 | 36 |
| C -> T | 224538 | SNP (transition) | 227 | 100.0% | 227 | 36 |
| C -> T | 543322 | SNP (transition) | 343 | 100.0% | 343 | 34 |
| G -> A | 553870 | SNP (transition) | 253 | 100.0% | 253 | 37 |
| T -> A | 635606 | SNP (transversion) | 284 | 100.0% | 284 | 36 |
| C -> T | 728478 | SNP (transition) | 293 | 100.0% | 293 | 35 |
| C -> T | 732272 | SNP (transition) | 290 | 100.0% | 290 | 34 |
| C -> T | 761836 | SNP (transition) | 242 | 100.0% | 242 | 35 |
| C -> T | 890575 | SNP (transition) | 267 | 100.0% | 267 | 36 |
| C -> T | 892510 | SNP (transition) | 287 | 100.0% | 287 | 35 |
| C -> T | 907987 | SNP (transition) | 236 | 99.6% | 235 | 36 |
| C -> T | 926164 | SNP (transition) | 143 | 100.0% | 143 | 35 |
| C -> T | 998214 | SNP (transition) | 279 | 100.0% | 279 | 36 |
| C -> T | 1028027 | SNP (transition) | 256 | 100.0% | 256 | 36 |
| C -> T | 1038116 | SNP (transition) | 287 | 100.0% | 287 | 35 |
| C -> T | 1226178 | SNP (transition) | 227 | 100.0% | 227 | 35 |
| G -> A | 1266744 | SNP (transition) | 231 | 100.0% | 231 | 36 |
| C -> T | 1280269 | SNP (transition) | 227 | 100.0% | 227 | 35 |
| C -> T | 1309222 | SNP (transition) | 282 | 100.0% | 282 | 36 |
| C -> T | 1337368 | SNP (transition) | 242 | 99.6% | 241 | 35 |
| C -> T | 1452465 | SNP (transition) | 222 | 99.5% | 221 | 34 |
| C -> T | 1485410 | SNP (transition) | 237 | 100.0% | 237 | 35 |
| C -> T | 1506876 | SNP (transition) | 252 | 99.6% | 251 | 36 |
| C -> T | 1525652 | SNP (transition) | 268 | 100.0% | 268 | 34 |
| C -> T | 1543574 | SNP (transition) | 282 | 100.0% | 282 | 35 |
| C -> T | 1579476 | SNP (transition) | 257 | 100.0% | 257 | 35 |
| G -> A | 1639122 | SNP (transition) | 269 | 100.0% | 269 | 35 |
| C -> T | 1678387 | SNP (transition) | 126 | 100.0% | 126 | 35 |
| C -> T | 1678389 | SNP (transition) | 123 | 100.0% | 123 | 35 |
| C -> T | 1726266 | SNP (transition) | 257 | 100.0% | 257 | 35 |
| G -> A | 1756092 | SNP (transition) | 199 | 99.5% | 198 | 35 |
| G -> A | 1928515 | SNP (transition) | 270 | 100.0% | 270 | 36 |
| G -> T | 1932209 | SNP (transversion) | 272 | 100.0% | 272 | 35 |
| G -> A | 1970577 | SNP (transition) | 342 | 99.4% | 340 | 35 |
| G -> A | 2251231 | SNP (transition) | 265 | 100.0% | 265 | 35 |
| G -> A | 2388218 | SNP (transition) | 263 | 100.0% | 263 | 36 |
| G -> A | 2389098 | SNP (transition) | 319 | 100.0% | 319 | 35 |
| C -> T | 2392562 | SNP (transition) | 147 | 100.0% | 147 | 34 |
| T -> C | 2411272 | SNP (transition) | 276 | 100.0% | 276 | 36 |
| G -> A | 2729902 | SNP (transition) | 298 | 100.0% | 298 | 35 |
| G -> A | 2878360 | SNP (transition) | 144 | 100.0% | 144 | 35 |
| G -> A | 2878362 | SNP (transition) | 143 | 100.0% | 143 | 34 |
| G -> A | 3191869 | Substitution | 267 | 99.3% | 265 | 35 |
| G -> A | 3191870 | Substitution | 268 | 99.3% | 266 | 35 |
| G -> A | 3249620 | SNP (transition) | 221 | 100.0% | 221 | 35 |
| G -> A | 3367008 | SNP (transition) | 316 | 99.1% | 313 | 35 |
| G -> A | 3553432 | SNP (transition) | 353 | 100.0% | 353 | 36 |
| G -> A | 3668754 | SNP (transition) | 309 | 100.0% | 309 | 36 |
| G -> A | 3809426 | SNP (transition) | 264 | 100.0% | 264 | 35 |
| G -> A | 3995026 | SNP (transition) | 265 | 100.0% | 265 | 36 |
| G -> A | 4368682 | SNP (transition) | 261 | 99.6% | 260 | 33 |
| C -> T | 4515045 | SNP (transition) | 237 | 100.0% | 237 | 35 |
| C -> T | 4535260 | SNP (transition) | 296 | 99.7% | 295 | 35 |
| C -> T | 4553943 | SNP (transition) | 277 | 99.3% | 275 | 34 |
| C -> T | 4591801 | SNP (transition) | 267 | 99.6% | 266 | 36 |
| C -> T | 4654377 | SNP (transition) | 280 | 100.0% | 280 | 36 |
| (T)7 -><br>(T)6 | 4687421 | Deletion | 158 | 100.0% | 158 | 29 |
| C -> T | 4694988 | SNP (transition) | 217 | 100.0% | 217 | 36 |
| C -> T | 4871633 | SNP (transition) | 266 | 100.0% | 266 | 33 |
| **SG491 *sifA*, *sseJ*, *sseL* WISH-8 JN4** |  |  |  |  |  |  |
| C -> T | 16626 | SNP (transition) | 251 | 99.6% | 250 | 31 |
| C -> T | 99756 | SNP (transition) | 303 | 99.7% | 302 | 34 |
| C -> T | 246774 | SNP (transition) | 258 | 100.0% | 258 | 35 |
| C -> T | 274696 | SNP (transition) | 253 | 100.0% | 253 | 35 |
| C -> T | 539398 | SNP (transition) | 252 | 98.8% | 249 | 35 |
| T -> A | 635606 | SNP (transversion) | 274 | 100.0% | 274 | 36 |
| C -> T | 645575 | SNP (transition) | 238 | 100.0% | 238 | 34 |
| G -> A | 680672 | SNP (transition) | 259 | 99.2% | 257 | 35 |
| C -> T | 739393 | SNP (transition) | 214 | 100.0% | 214 | 35 |
| C -> T | 829063 | SNP (transition) | 190 | 100.0% | 190 | 34 |
| C -> T | 842242 | SNP (transition) | 213 | 98.1% | 209 | 35 |
| C -> T | 877122 | SNP (transition) | 266 | 100.0% | 266 | 30 |
| C -> T | 928145 | SNP (transition) | 244 | 100.0% | 244 | 35 |
| C -> T | 1034782 | SNP (transition) | 297 | 99.7% | 296 | 33 |
| C -> T | 1047125 | SNP (transition) | 260 | 100.0% | 260 | 35 |
| C -> T | 1262555 | SNP (transition) | 262 | 100.0% | 262 | 35 |
| G -> A | 1266744 | SNP (transition) | 211 | 100.0% | 211 | 35 |
| C -> T | 1326912 | SNP (transition) | 240 | 99.6% | 239 | 35 |
| C -> T | 1460270 | SNP (transition) | 223 | 100.0% | 223 | 35 |
| C -> T | 1673248 | SNP (transition) | 236 | 99.6% | 235 | 36 |
| C -> T | 1678387 | SNP (transition) | 200 | 99.5% | 199 | 35 |
| C -> T | 1678389 | SNP (transition) | 199 | 100.0% | 199 | 34 |
| G -> A | 1703437 | SNP (transition) | 201 | 100.0% | 201 | 34 |
| G -> A | 1711108 | SNP (transition) | 224 | 100.0% | 224 | 36 |
| G -> A | 1719692 | SNP (transition) | 258 | 99.6% | 257 | 35 |
| G -> A | 1737233 | SNP (transition) | 228 | 99.6% | 227 | 35 |
| G -> A | 1748229 | SNP (transition) | 201 | 100.0% | 201 | 36 |
| G -> A | 1756092 | SNP (transition) | 211 | 100.0% | 211 | 35 |
| G -> A | 1801868 | SNP (transition) | 238 | 100.0% | 238 | 35 |
| G -> A | 1812321 | SNP (transition) | 224 | 99.6% | 223 | 34 |
| G -> A | 1822302 | SNP (transition) | 215 | 99.5% | 214 | 35 |
| G -> A | 1844257 | SNP (transition) | 258 | 100.0% | 258 | 33 |
| G -> A | 1852123 | SNP (transition) | 268 | 100.0% | 268 | 35 |
| G -> A | 1880959 | SNP (transition) | 245 | 100.0% | 245 | 36 |
| G -> A | 1893905 | SNP (transition) | 257 | 99.6% | 256 | 35 |
| G -> A | 1910319 | SNP (transition) | 208 | 100.0% | 208 | 36 |
| G -> A | 1956581 | SNP (transition) | 221 | 100.0% | 221 | 35 |
| G -> A | 1961994 | SNP (transition) | 241 | 100.0% | 241 | 35 |
| G -> A | 1984340 | SNP (transition) | 175 | 100.0% | 175 | 37 |
| G -> A | 2010519 | SNP (transition) | 278 | 100.0% | 278 | 35 |
| G -> A | 2051044 | SNP (transition) | 237 | 100.0% | 237 | 36 |
| G -> A | 2154098 | SNP (transition) | 263 | 99.6% | 262 | 35 |
| G -> A | 2274736 | SNP (transition) | 217 | 100.0% | 217 | 34 |
| G -> A | 2277779 | SNP (transition) | 229 | 100.0% | 229 | 36 |
| G -> A | 2292499 | SNP (transition) | 243 | 100.0% | 243 | 35 |
| G -> A | 2320931 | SNP (transition) | 279 | 100.0% | 279 | 35 |
| G -> A | 2336076 | SNP (transition) | 201 | 100.0% | 201 | 35 |
| C -> T | 2392562 | SNP (transition) | 214 | 100.0% | 214 | 34 |
| T -> C | 2411272 | SNP (transition) | 290 | 100.0% | 290 | 37 |
| G -> A | 2493849 | SNP (transition) | 251 | 100.0% | 251 | 37 |
| G -> A | 2537197 | SNP (transition) | 246 | 99.6% | 245 | 36 |
| G -> A | 2608719 | SNP (transition) | 272 | 99.6% | 271 | 35 |
| G -> A | 2618227 | SNP (transition) | 228 | 100.0% | 228 | 34 |
| G -> A | 2802657 | SNP (transition) | 227 | 100.0% | 227 | 36 |
| G -> A | 2841282 | SNP (transition) | 290 | 100.0% | 290 | 35 |
| G -> A | 2867053 | SNP (transition) | 257 | 100.0% | 257 | 35 |
| G -> A | 3004903 | SNP (transition) | 226 | 100.0% | 226 | 35 |
| G -> A | 3004981 | SNP (transition) | 255 | 100.0% | 255 | 34 |
| G -> A | 3075993 | SNP (transition) | 285 | 99.6% | 284 | 35 |
| G -> A | 3440034 | SNP (transition) | 234 | 100.0% | 234 | 34 |
| G -> A | 3441452 | SNP (transition) | 298 | 100.0% | 298 | 36 |
| G -> A | 3451254 | SNP (transition) | 326 | 100.0% | 326 | 35 |
| G -> A | 3675197 | SNP (transition) | 314 | 100.0% | 314 | 36 |
| G -> A | 3695408 | SNP (transition) | 311 | 99.4% | 309 | 34 |
| C -> T | 3841824 | SNP (transition) | 276 | 100.0% | 276 | 35 |
| G -> A | 3843533 | SNP (transition) | 265 | 100.0% | 265 | 34 |
| G -> A | 4104124 | SNP (transition) | 221 | 99.5% | 220 | 32 |
| C -> T | 4129199 | SNP (transition) | 256 | 100.0% | 256 | 36 |
| C -> T | 4145068 | SNP (transition) | 275 | 99.6% | 274 | 35 |
| C -> T | 4157626 | SNP (transition) | 229 | 100.0% | 229 | 35 |
| C -> T | 4166821 | SNP (transition) | 295 | 99.0% | 292 | 35 |
| C -> T | 4287312 | SNP (transition) | 263 | 100.0% | 263 | 35 |
| G -> A | 4368682 | SNP (transition) | 244 | 100.0% | 244 | 33 |
| C -> T | 4569786 | SNP (transition) | 227 | 100.0% | 227 | 36 |
| **SG457 *sifA*, *sseJ*, *sseL* WISH-10 JN7** |  |  |  |  |  |  |
| C -> T | 169880 | SNP (transition) | 263 | 100.0% | 263 | 36 |
| T -> A | 635606 | SNP (transversion) | 245 | 99.6% | 244 | 37 |
| C -> T | 743345 | SNP (transition) | 189 | 100.0% | 189 | 35 |
| C -> T | 1131356 | SNP (transition) | 221 | 100.0% | 221 | 36 |
| G -> A | 1266744 | SNP (transition) | 205 | 100.0% | 205 | 36 |
| C -> T | 1289757 | SNP (transition) | 198 | 100.0% | 198 | 36 |
| C -> T | 1500959 | SNP (transition) | 200 | 100.0% | 200 | 36 |
| C -> T | 1678384 | SNP (transition) | 202 | 100.0% | 202 | 36 |
| C -> T | 1678387 | SNP (transition) | 201 | 100.0% | 201 | 34 |
| C -> T | 1678389 | SNP (transition) | 201 | 100.0% | 201 | 35 |
| G -> A | 1756092 | SNP (transition) | 226 | 100.0% | 226 | 35 |
| G -> A | 1782243 | SNP (transition) | 183 | 99.5% | 182 | 36 |
| G -> A | 2025576 | SNP (transition) | 214 | 100.0% | 214 | 36 |
| G -> A | 2222977 | SNP (transition) | 192 | 99.5% | 191 | 35 |
| C -> T | 2392562 | SNP (transition) | 174 | 100.0% | 174 | 34 |
| T -> C | 2411272 | SNP (transition) | 238 | 100.0% | 238 | 37 |
| C -> T | 2829893 | SNP (transition) | 268 | 99.6% | 267 | 35 |
| G -> A | 2980682 | SNP (transition) | 192 | 100.0% | 192 | 36 |
| C -> T | 3460714 | SNP (transition) | 263 | 98.9% | 260 | 34 |
| G -> A | 3794301 | SNP (transition) | 243 | 99.6% | 242 | 35 |
| G -> A | 3795782 | SNP (transition) | 219 | 99.5% | 218 | 36 |
| G -> A | 3833844 | SNP (transition) | 236 | 100.0% | 236 | 34 |
| G -> A | 3838846 | SNP (transition) | 196 | 100.0% | 196 | 36 |
| G -> A | 3964681 | SNP (transition) | 281 | 100.0% | 281 | 28 |
| G -> A | 4027923 | SNP (transition) | 238 | 99.6% | 237 | 35 |
| C -> T | 4298145 | SNP (transition) | 215 | 100.0% | 215 | 35 |
| G -> A | 4368682 | SNP (transition) | 247 | 99.6% | 246 | 34 |
| C -> T | 4828236 | SNP (transition) | 241 | 100.0% | 241 | 35 |

